# Regulation of Neuroendocrine Plasticity by the RNA-Binding Protein ZFP36L1

**DOI:** 10.1101/2021.10.22.465501

**Authors:** Hsiao-Yun Chen, Yavuz T. Durmaz, Yixiang Li, Amin H. Sabet, Amir Vajdi, Thomas Denize, Emily Walton, John G. Doench, Navin R. Mahadevan, Julie-Aurore Losman, David A. Barbie, Michael Y. Tolstorukov, Charles M. Rudin, Triparna Sen, Sabina Signoretti, Matthew G. Oser

## Abstract

Some small cell lung cancers (SCLCs) are highly sensitive to inhibitors of the histone demethylase LSD1. LSD1 inhibitors are thought to induce their anti-proliferative effects by blocking neuroendocrine differentiation, but the mechanisms by which LSD1 controls the SCLC neuroendocrine phenotype are not well understood. To identify genes required for LSD1 inhibitor sensitivity in SCLC, we performed a positive selection genome-wide CRISPR/Cas9 loss of function screen and found that *ZFP36L1*, an mRNA-binding protein that destabilizes mRNAs, is required for LSD1 inhibitor sensitivity. LSD1 binds and represses ZFP36L1 and upon LSD1 inhibition, ZFP36L1 expression is restored, which is sufficient to block the SCLC neuroendocrine differentiation phenotype and induce a non-neuroendocrine “inflammatory” phenotype. Mechanistically, ZFP36L1 binds and destabilizes SOX2 and INSM1 mRNAs, two transcription factors that are required for SCLC neuroendocrine differentiation. This work identifies ZFP36L1 as an LSD1 target gene that controls the SCLC neuroendocrine phenotype and demonstrates that modulating mRNA stability of lineage transcription factors controls neuroendocrine to non-neuroendocrine plasticity.

## Introduction

Small cell lung cancer (SCLC) is a high-grade neuroendocrine cancer that accounts for 15% of lung cancers^1–3^. Most SCLC patients initially respond to conventional chemotherapy, but nearly all experience a recurrence of the disease for which there are currently no effective treatment options. SCLCs are driven by near universal loss of function (LOF) mutations in tumor suppressor genes *RB1* and *TP53* and do not harbor recurrent mutations in a druggable oncogenic driver^4–6^, which makes finding therapeutic targets challenging. There are no approved targeted therapies for SCLC.

SCLC and acute myeloid leukemia (AML) cell lines are selectively sensitive to inhibition of the histone demethylase LSD1, while nearly all other cancer cell lines are inherently resistant^7, 8^. This has inspired development of several LSD1 inhibitors for clinical trials in SCLC and AML patients^7–11^. Preclinical studies in SCLC cell lines and patient-derived xenografts (PDXs) show that some SCLC models are highly sensitive to LSD1 inhibitors while others are more resistant^7, 10^. An understanding of the molecular basis for why some SCLCs are sensitive to LSD1 inhibitors could lead to the identification of biomarkers for patient selection.

More than 70% of SCLCs highly express the lineage specific neuroendocrine transcription factor ASCL1^3, 4, 12^ that functions to transcriptionally maintain SCLC in a neuroendocrine differentiation state, which is necessary for SCLC tumorigenesis and survival^13, 14^. LSD1 inhibition dramatically downregulates ASCL1 and neuroendocrine differentiation in some SCLC cell lines and ASCL1 downregulation correlates with LSD1 inhibitor sensitivity in SCLC cell lines and PDX models^7^. Similarly, inhibition of KDM5A or the KDM5-family of histone demethylases also represses ASCL1 expression^15^. Moreover, combined inhibition of LSD1 and KDM5 histone demethylases synergistically represses ASCL1 and SCLC proliferation^15^. LSD1 or KDM5A inhibition upregulates NOTCH and suppresses ASCL1 through a NOTCH-dependent mechanism^7, 15^. Both LSD1 and KDM5A are H3K4 histone demethylases that canonically function to repress target genes, but have distinct substrate specificities. LSD1 preferentially demethylates mono- or di-methylated lysine 4 on histone 3 (H3K4) at enhancers, while KDM5A preferentially demethylates tri-methylated H3K4 at promoters^16–20^. Therefore, LSD1 and KDM5A could function in a cooperative manner to repress NOTCH and sustain the SCLC neuroendocrine phenotype. However, apart from LSD1’s and KDM5A’s ability to repress NOTCH, it is not known why only some SCLCs are sensitive to LSD1 inhibition and how LSD1 and KDM5A function to promote SCLC neuroendocrine differentiation and survival.

Herein, we performed a genome-wide CRISPR/Cas9 positive selection screen in a SCLC cell line treated with an LSD1 inhibitor alone or combined with a KDM5 inhibitor. Our screen systematically uncovered mechanisms required for LSD1 inhibitor sensitivity and discovered critical target genes repressed by LSD1 that are required for the ASCL1-driven SCLC neuroendocrine state and survival.

## Results

### CRISPR/Cas9 Positive Selection Screen Identifies Genes Required for LSD1 Inhibitor Sensitivity in Small Cell Lung Cancer

Combined LSD1 and KDM5 inhibition with the small molecular inhibitors ORY-1001 and KDM5-C70, respectively, synergistically blocks ASCL1 and cellular proliferation in some SCLC cell lines, including the NCI-H1876 SCLC cell line where ORY-1001 combined with KDM5-C70 nearly completely represses ASCL1 and cellular proliferation^15^. To systematically identify genes required for LSD1 and KDM5 inhibitor sensitivity, we performed a CRISPR/Cas9 positive selection knockout screen in the presence of ORY-1001 alone or ORY-1001 combined with KDM5-C70. Given the strong positive correlation between decreased ASCL1 expression and decreased proliferation upon LSD1 and/or KDM5 inhibitor treatment^7, 15^, we hypothesized that our screen would also uncover genes that regulate ASCL1 and neuroendocrine differentiation in SCLC.

NCI-H1876 cells were engineered to express Cas9 and then infected with the genome-wide Brunello sgRNA library at an MOI of ∼0.3 maintaining representation of >500 cells/guide for each arm of the screen. Following puromycin selection and allowing time for maximal CRISPR/Cas9 editing, puromycin-resistant cells were then split into 4 arms with each arm treated with one of two low concentrations of the LSD1 inhibitor ORY-1001 (1 or 100 nM), alone or combined with the KDM5 inhibitor, KDM5-C70 (1000 nM) (Fig. 1A). After 24 days when there was selective growth of sgRNA-infected cells compared to mock treated cells in the presence of drug (Fig. 1B), the cells were harvested and their genomic DNA was isolated and sent for deep sequencing. Analysis of sgRNA enrichment at the end time point compared to the initial timepoint showed striking enrichment of some sgRNAs in both biological replicates (Extended Data Fig. S1A-D). Hypergeometric and STARS analysis revealed several highly statistically significant hits in the ORY-1001 and the ORY-1001/KDM5-C70 treatment arms (Fig. 1C-F and Extended Data Fig. S1E-H; Extended Data Table S1). Our expected positive control *KMT2C,* an H3K4 methyltransferase whose enzymatic activity opposes the catalytic activity of LSD1 at enhancers, scored as one of the top hits in the ORY-1001 screen (Extended Data Fig. S1E) indicating that our screen was effective. A top hit in the combined treatment arm was REST (Extended Data Fig. S1G,H), a transcriptional repressor of neuronal genes^21^. REST is an LSD1- target gene^7, 22^ and a NOTCH-target gene that is necessary and sufficient to induce a non-neuroendocrine subpopulation of cells during SCLC tumorigenesis^22, 23^. These results are consistent with previous data demonstrating that LSD1 and KDM5 inhibitors mediate their anti-proliferative effects through a NOTCH-dependent mechanism^7, 15^ and suggest that the NOTCH-dependent mechanism converges on REST. Our screen also identified several novel hits that were not previously linked to LSD1 or SCLC neuroendocrine differentiation (Fig. 1C-F and Extended Data Fig. S1E-H). For example, the RNA-binding protein *ZFP36L1* scored as a top 5 hit in all arms of the screen and has not been previously linked to LSD1 or KDM5 or studied in SCLC. Other hits were selectively enriched in the combined treatment arms and were comprised of genes within the same pathway including *YAP1* and *WWTR1* (TAZ); and *RUNX1* and its binding partner *CBFB* (Fig. 1E,F and Extended Data Fig. S1G,H). Lastly, LZTR1, a regulator of RAS protein stability, selectively scored in the ORY-1001 arm (Fig. 1C, D and Extended Data Fig. S1E,F). Together, these findings suggest that our screen was robust and uncovered several candidate genes required for sensitivity to LSD1 inhibition alone or combined LSD1/KDM5 inhibition.

**Fig. 1:**
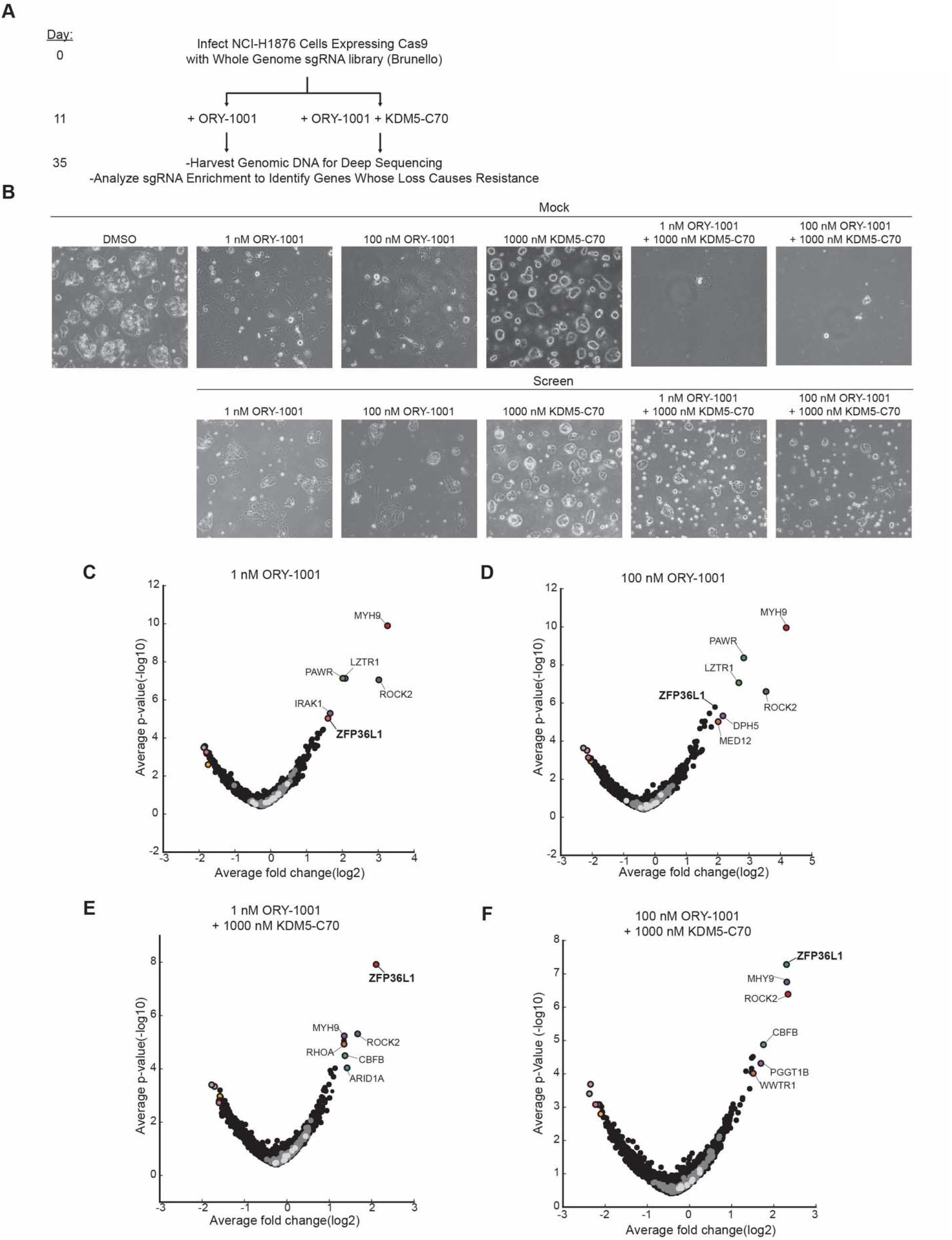
CRISPR/Cas9 Positive Selection Screen Identifies Genes Required for LSD1 Inhibitor Sensitivity in Small Cell Lung Cancer. (**A**)Schema for the CRISPR/Cas9 positive selection screen in NCI-H1876 Cas9 cells infected with the Brunello sgRNA library. Following selection, cells were treated with one of two low concentrations of the LSD1 inhibitor ORY-1001 (1 or 100 nM) alone or combined with the KDM5 inhibitor, KDM5-C70 (1000 nM), and then harvested at day 35 when the sgRNA-infected cells showed resistance. (**B**) Representative brightfield images of NCI-H1876 Cas9 cells infected with the Brunello sgRNA library or mock-infected at the day 35 screen endpoint. (**C-F**) Hypergeometric analysis from the positive-selection CRISPR/Cas9 screen on day 35 relative to the early timepoint prior to drug treatment (day 11) of NCI-H1876 Cas9 cells infected with the Brunello sgRNA library and then treated with ORY-1001 (1 nM) (**C**), ORY-1001 (100 nM) (**D**), ORY-1001 (1 nM) + KDM5-C70 (1000 nM) (**E**), or ORY-1001 (100 nM) + KDM5-C70 (1000 nM) (**F**). n=2 biological replicates.

### ZFP36L1, REST, YAP1, and WWTR1, are LSD1 Target Genes De-Repressed by LSD1 Inhibition

For further study, we focused on hits that were most highly statistically significant, hits that scored highly in multiple arms of our screen, and hits where multiple genes within a pathway scored. We reasoned that the hits would segregate into 2 broad categories: anti-proliferative LSD1 target genes normally repressed by LSD1 and de-repressed by ORY-1001; and genes that regulate parallel pathways that render the cell insensitive to LSD1 inhibition, but whose mRNA expression is unchanged by ORY-1001. To distinguish these possibilities, we performed RT-qPCR in NCI-H1876 cells after ORY-1001 or ORY-1001/KDM5-C70 treatment. The mRNAs of several hits including *ZFP36L1*, *REST* [a known LSD1 target gene^7, 22^], and *YAP1* were significantly upregulated after ORY-1001 treatment (Fig. 2A) suggesting that these hits are LSD1 target genes de-repressed upon ORY-1001 treatment and required for ORY- 1001’s anti-proliferative response. In contrast, ORY-1001 did not alter mRNA expression of *LZTR1*, *CBFB*, and *RUNX1* indicating that these genes likely caused resistance by modulating the activity of parallel pathways required for LSD1 inhibitor sensitivity. Combined treatment with ORY-1001 and KDM5-C70 did not further upregulate mRNA expression of *ZFP36L1*, *REST*, *YAP1*, and *WWTR1.* Therefore, subsequent validation efforts focused on LSD1 inhibition alone.

**Fig. 2:**
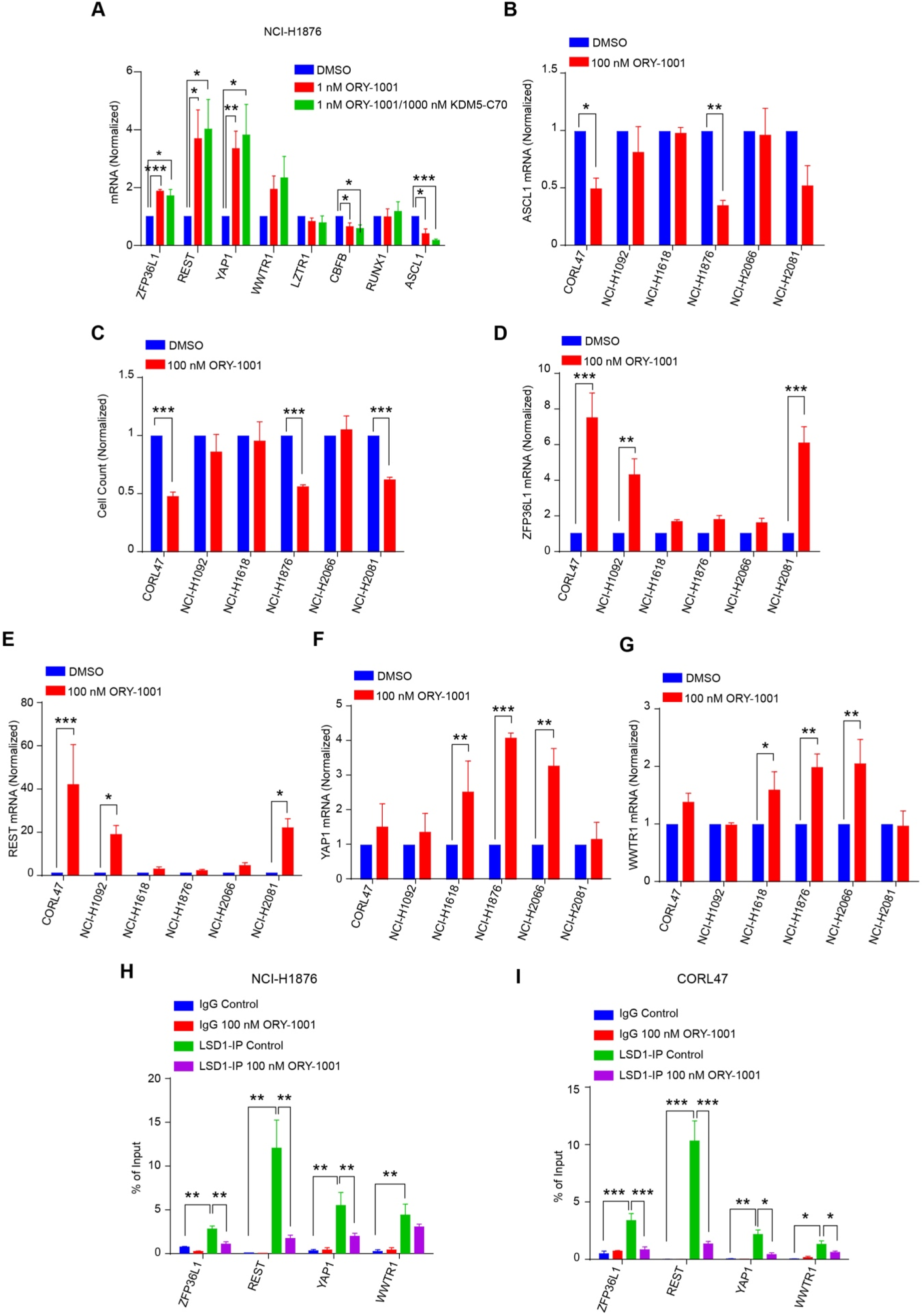
ZFP36L1, REST, YAP1, and WWTR1 are Bound by LSD1 and De-Repressed by LSD1 Inhibition. (**A**) RT-qPCR of NCI-H1876 cells treated with ORY-1001 (1 nM) or ORY-1001 (1 nM) + KDM5- C70 (1000 nM) for 6 days. n=4 biological replicates. (**B and D-G**) RT-qPCR for *ASCL1* (**B**), *ZFP36L1* (**D**), *REST* (**E**), *YAP1* (**F**), and *WWTR1* (**G**) in different human SCLC cell lines treated with ORY-1001 (100 nM) or DMSO for 6 days. For B, n=3 biological replicates. For D-G, n=2 biological replicates. (**C**) Cellular proliferation of human SCLC cell lines treated with ORY-1001 (100 nM) for 6 days relative to the cells treated with DMSO. n=3 biological replicates. (**H and I**) ChIP-qPCR of NCI-H1876 (**H**) and CORL47 (**I**) cells treated with ORY-1001 (100 nM) or DMSO for 6 days followed by immunoprecipitation (IP) for LSD1 and qPCR for the genes indicated. See S2A-D for the identification of enrichment peaks in the genes where LSD1 is bound. n=2 biological replicates. For all panels, *=p<0.05, **=p<0.01, ***=p<0.001.

We then asked whether ORY-1001 treatment induces *ZFP36L1*, *REST*, *YAP1*, and *WWTR1* mRNA expression across multiple ASCL1-positive SCLC cell lines. Consistent with previous findings^7^, ORY-1001 repression of ASCL1 correlated with ORY-1001 sensitivity (Fig. 2B,C). ORY-1001 induced *ZFP36L1*, *REST*, *YAP1*, and *WWTR1* expression across multiple SCLC cell lines (Fig. 2D-G). The magnitude of induction for each gene varied among cell lines. *ZFP36L1* and *REST* were most strongly induced in CORL47, NCI-H1092, and NCI-H2081 cells (Fig. 2D,E); whereas Y*AP1* and *WWTR1* were more selectively induced in NCI-H1618, NCI- H1876, and NCI-H2066 cells (Fig. 2F,G).

To test whether LSD1 binds *ZFP36L1*, *REST*, *YAP1*, and *WWTR1* directly, we performed LSD1 chromatin immunoprecipitation (ChIP) followed by qPCR (ChIP-qPCR) in NCI- H1876 and CORL47 cells. LSD1 binding regions in *ZFP36L1*, *REST*, *YAP1*, and *WWTR1* were identified using LSD1, H3K4me1, and H3K4me2 ChIP-Sequencing data from cistrome.db^24, 25^ (Extended Data Fig. S2A-D). LSD1 inhibitors can displace LSD1 from binding chromatin ^26^ and therefore ChIP-qPCR was performed in the presence or absence of ORY-1001. Compared to the IgG isotype-matched control, LSD1 bound to *ZFP36L1*, *REST*, *YAP1*, and *WWTR1* in untreated CORL47 and NCI-H1876 cells, and binding was blocked by ORY-1001 (Fig. 2H-I and Extended Data Fig. S2E,F). Collectively, our data demonstrates that LSD1 binds *ZFP36L1*, *YAP1*, *WWTR1*, and *REST,* and that LSD1 inhibition displaces LSD1 from binding these targets and induces their mRNA expression.

### ZFP36L1 is Repressed in Small Cell Lung Cancer

We then focused on *ZFP36L1* as a novel LSD1 target gene (see Fig. 2) that is required for LSD1 inhibitor sensitivity (see Fig. 1). ZFP36L1 is an mRNA-binding protein that binds AUUUA/UUAUUUAUU motifs in the 3’ UTR of target genes and degrades their mRNA^27^. Previous work has demonstrated that ZFP36L1 promotes cell quiescence^28^ and functions as a tumor suppressor in acute lymphoblastic leukemia (ALL)^29^. Interestingly, ZFP36L1 mRNA expression is only repressed in 3 cancer types: SCLC; Neuroblastoma, a pediatric neural crest tumor that like SCLC is ASCL1-dependent^30^; and AML, which like SCLC, is uniquely sensitive to LSD1 inhibitors^7–9^ (Fig. 3A). Consistent with the SCLC cell line data, RNA-sequencing from human SCLC tumors shows that *ZFP36L1* mRNA expression is markedly reduced in SCLC compared to lung adenocarcinoma (Fig. 3B). This was confirmed at the protein level by immunohistochemistry (IHC) for ZFP36L1 using human lung tumor tissue microarrays (TMAs) (Fig. 3C,D). Zfp36l1 protein expression was also absent in the majority of cells in mouse SCLC lung tumors derived from a CRISPR-based SCLC genetically-engineered mouse model^15^ (Fig. 3E, F). Interestingly, there are subpopulations of cells within these mouse SCLC tumors where Zfp36l1 is expressed, and these subpopulations have relatively lower expression of Ascl1 (Fig. 3G) suggesting a negative correlation between Zfp36l1 and Ascl1 expression.

**Fig. 3:**
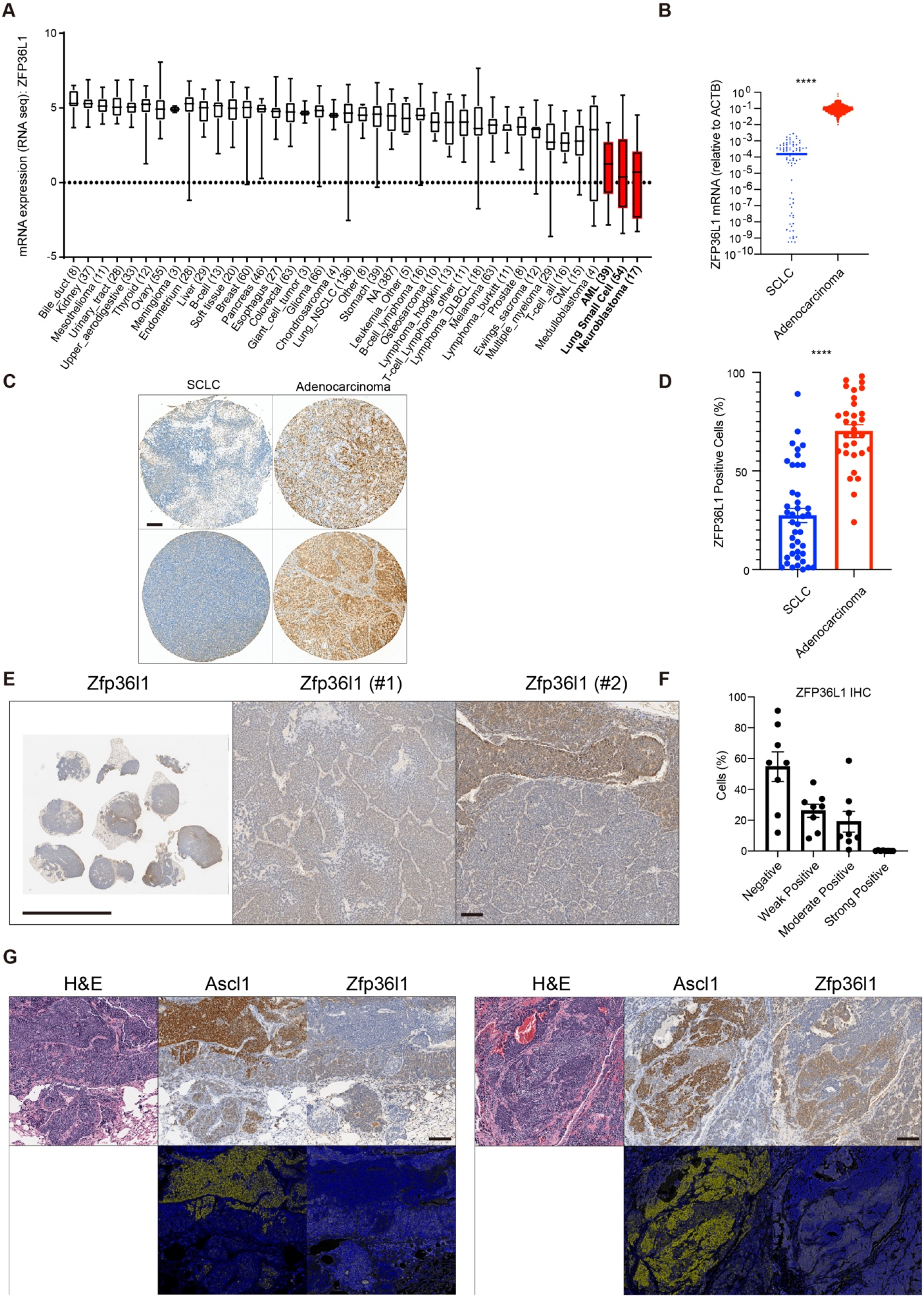
ZFP36L1 is Repressed in Small Cell Lung Cancer. (**A**) ZFP36L1 mRNA expression in cell lines from the cancer cell line encyclopedia (CCLE) RNA-sequencing dataset. AML, SCLC, and Neuroblastoma are highlighted in red. (**B**) ZFP36L1 mRNA expression in human SCLC and lung adenocarcinoma tumors normalized to *ACTB* from publicly available RNA-sequencing data on cbioportal.org^49^. n=81 SCLC tumors, n=747 lung adenocarcinoma tumors. (**C**) Representative IHC for ZFP36L1 from human tissue microarrays (TMAs) of SCLC and lung adenocarcinoma. Bar=100 μM. (**D**) Quantitation of all SCLC and lung adenocarcinoma specimens on the TMAs. n=41 SCLC tumors, n=31 lung adenocarcinoma tumors. (**E**) Representative IHC for Zfp36l1 in mouse SCLC lung tumors from a CRISPR-based SCLC genetically-engineered mouse model. Scale Bar-left=1 cm, right=100 μm. (**F**) Quantitation of % positive Zfp36l1 cells in SCLC mouse tumors. n=8 lung tumors. (**G**) Pseudo-colored Zfp36l1 and Ascl1 IHC highlighting the negative correlation between Zfp36l1 and Ascl1. Bar=100 μM. For all panels, ****=p<0.0001.

### LSD1 Inhibitors Block Neuroendocrine Differentiation and Proliferation in Small Cell Lung Cancer Through a ZFP36L1-Dependent Mechanism

Time-course experiments after ORY-1001 in NCI-H1876, CORL47, and NCI-H69 cells confirmed that ZFP36L1 expression is markedly induced after 7 to 8 days of ORY-1001 treatment (Fig. 4A,B and Extended Data Fig. S3A-C). Similarly, treatment with the structurally unrelated non-covalent LSD1 inhibitor CC-90011^10^ induced ZFP36L1 expression and inhibited cellular proliferation (Extended Data Fig. S3D-F). Given that ZFP36L1 is: 1. Repressed in SCLC tumors and cell lines; 2. Expressed in lung adenocarcinoma and in subpopulations of SCLC tumors with lower ASCL1 expression; and 3. Induced by LSD1 inhibitor treatment while ASCL1 is repressed, we hypothesized that LSD1 repression of ZFP36L1 is required to sustain ASCL1 and SCLC neuroendocrine differentiation. To test this, we used CRISPR/Cas9 to generate NCI- H1876 and CORL47 cells that were either ZFP36L1-knockout (sgZFP36L1) or ZFP36L1-WT (sgControl) and performed studies with ORY-1001 or CC-90011. Both ORY-1001 and CC- 90011 decreased ASCL1 mRNA expression and protein levels and blocked cellular proliferation in NCI-H1876 and CORL47 sgControl cells (Fig. 4C-E and Extended Data Fig. S4A-I). These effects were significantly attenuated in NCI-H1876 and CORL47 sgZFP36L1 cells (Fig. 4C-E and Extended Data Fig. S4A-I and S5A-B), demonstrating that ZFP36L1 plays an important role in the repression of ASCL1 and cellular proliferation by LSD1 inhibitors.

**Fig. 4:**
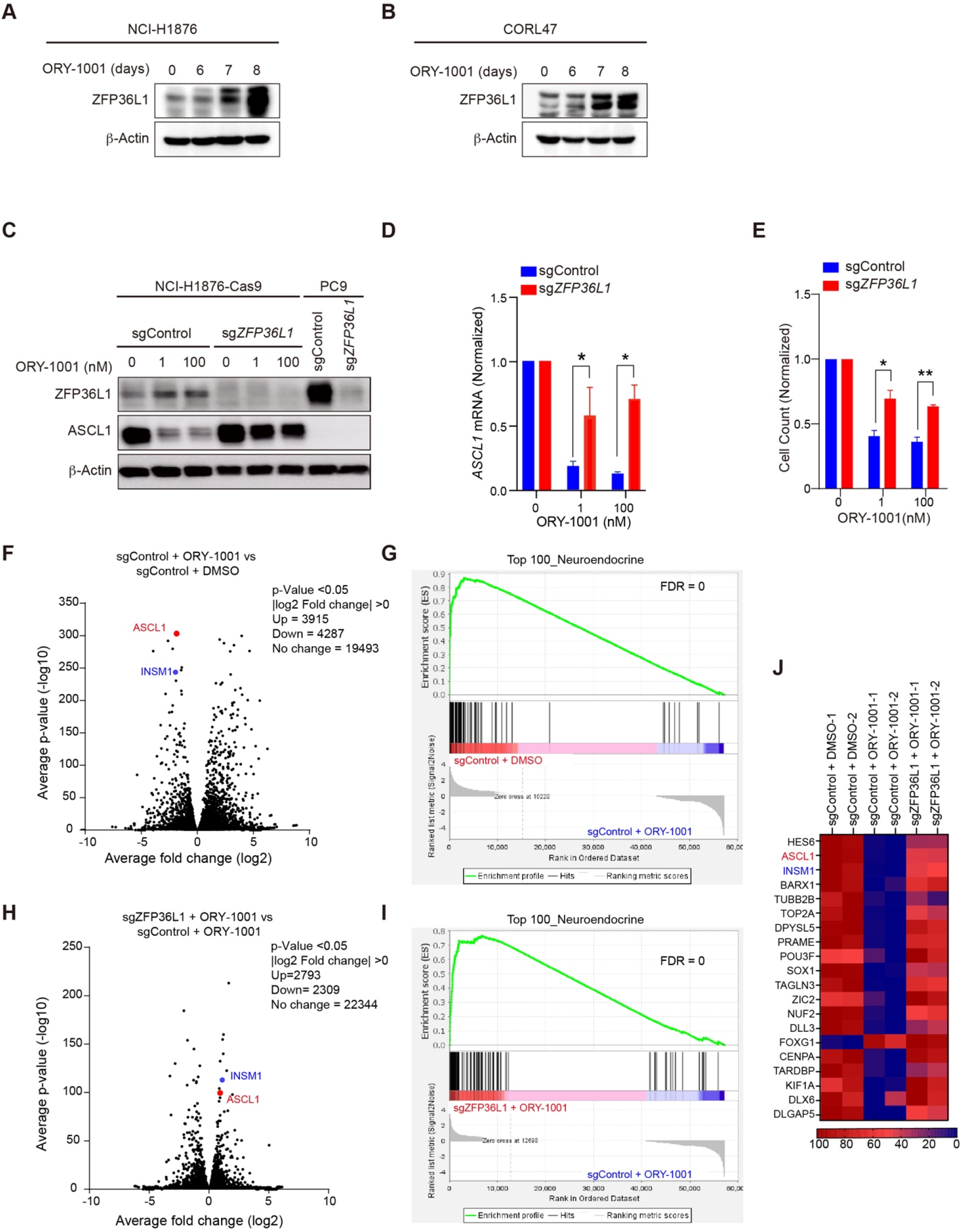
LSD1 Inhibitors Block Neuroendocrine Differentiation and Proliferation in Small Cell Lung Cancer Through a ZFP36L1-Dependent Mechanism. (**A and B**) Immunoblot analysis of NCI-H1876 cells (**A**) and CORL47 cells (**B**) treated with ORY- 1001 (100 nM) for 6, 7 and 8 days. (**C-E**) Immunoblot analysis (**C**), RT-qPCR (**D**), and quantitation of cell counts (**E**) of NCI-H1876 Cas9 cells infected lentiviruses encoding an sgRNA targeting ZFP36L1 (sgZFP36L1) or a non-targeting sgRNA (sgControl) and then treated with ORY-1001 (1 nM and 100 nM) or DMSO for 7 days. For D, n=2 biological replicates. For E, n=3 biological replicates. (**F and H**) Volcano plot of RNA-sequencing (RNA-seq) data showing the log2 fold change and -Log10 p-value of gene expression comparing NCI-H1876 sgControl cells treated with ORY-1001 (1 nM) or DMSO (**F**) or comparing NCI-H1876 sgZFP36L1 cells treated with ORY-1001 (1 nM) vs. H1876 sgControl cells treated with ORY-1001 (1 nM) (**H**). ASCL1 and INSM1 are indicated in red and blue, respectively. n=2 biological replicates. (**G and I**) Gene set enrichment analysis (GSEA) of RNA-seq data in F (**G**) or H (**I**) of the Top 100 Neuroendocrine genes. FDR q-values are indicated. (**J**) Heatmap from the RNA-seq experiment in F-I of the cells indicated of the top 20 Neuroendocrine Genes from the Top 100 Neuroendocrine gene set. Red denotes genes with high expression, and blue denotes genes with low expression. For all panels, *=p<0.05, **=p<0.01.

We then used NCI-H1876 ZFP36L1-knockout or ZFP36L1-WT cells treated with ORY- 1001 or DMSO to perform RNA-sequencing (Extended Data Fig. S6) followed by gene set enrichment analysis (GSEA). ORY-1001 treatment of sgControl cells dramatically reduced neuroendocrine genes and ASCL1 target genes (Fig. 4F,G,J and Extended Data Fig. S7), which is consistent with a previous study^7^. ZFP36L1 knockout markedly rescued the loss of neuroendocrine genes and ASCL1 target genes caused by ORY-1001 (Fig. 4H-J and Extended Data Fig. S7). These results demonstrate that ZFP36L1 is required for LSD1 inhibitors to block neuroendocrine differentiation.

### ZFP36L1 Repression is Required for SCLC Proliferation and Neuroendocrine Differentiation

We then asked if ZFP36L1 expression is sufficient to block neuroendocrine differentiation and cellular proliferation in SCLC. To activate the endogenous ZFP36L1 locus, we co-expressed a deactivated Cas9 fused to the transcriptional activator VP64 (dCas9-VP64)^31^ and sgRNAs targeting the ZFP36L1 promoter in CORL47 cells. Two independent sgRNAs targeting the ZFP36L1 promoter induced ZFP36L1 mRNA expression and protein expression in CORL47 cells (Fig. 5A,B) and decreased protein levels of ASCL1 and SOX2, another lineage specific transcription factor required for SCLC survival and lineage plasticity in neuroendocrine prostate cancer^6, 32^ (Fig. 5B). This correlated with a decrease in cellular proliferation (Fig. 5C). Similarly, exogenous expression of ZFP36L1 decreased ASCL1 and SOX2, and markedly decreased cellular proliferation in CORL47 and NCI-H69 cells (Fig. 5E,F and Extended Data Fig. S8A,B). We then engineered a ZFP36L1-mRNA binding mutant where 8 cysteines or histidines are mutated to alanines, which was previously shown to block the ability of ZFP36L1 to bind mRNA^33^ (Fig. 5D), and expressed the ZFP36L1-mRNA-binding mutant or ZFP36L1-WT constitutively or inducibly (using a DOX-ON system) in CORL47 cells. Re-expression of ZFP36L1 WT, but not the ZFP36L1-mRNA-binding mutant, decreased neuroendocrine differentiation and cellular proliferation (Fig. 5E-H and Extended Data Fig. S8A,B). This demonstrated that ZFP36L1 expression is sufficient to block neuroendocrine differentiation and cellular proliferation in SCLC and that these phenotypes require ZFP36L1’s mRNA binding activity.

**Fig. 5:**
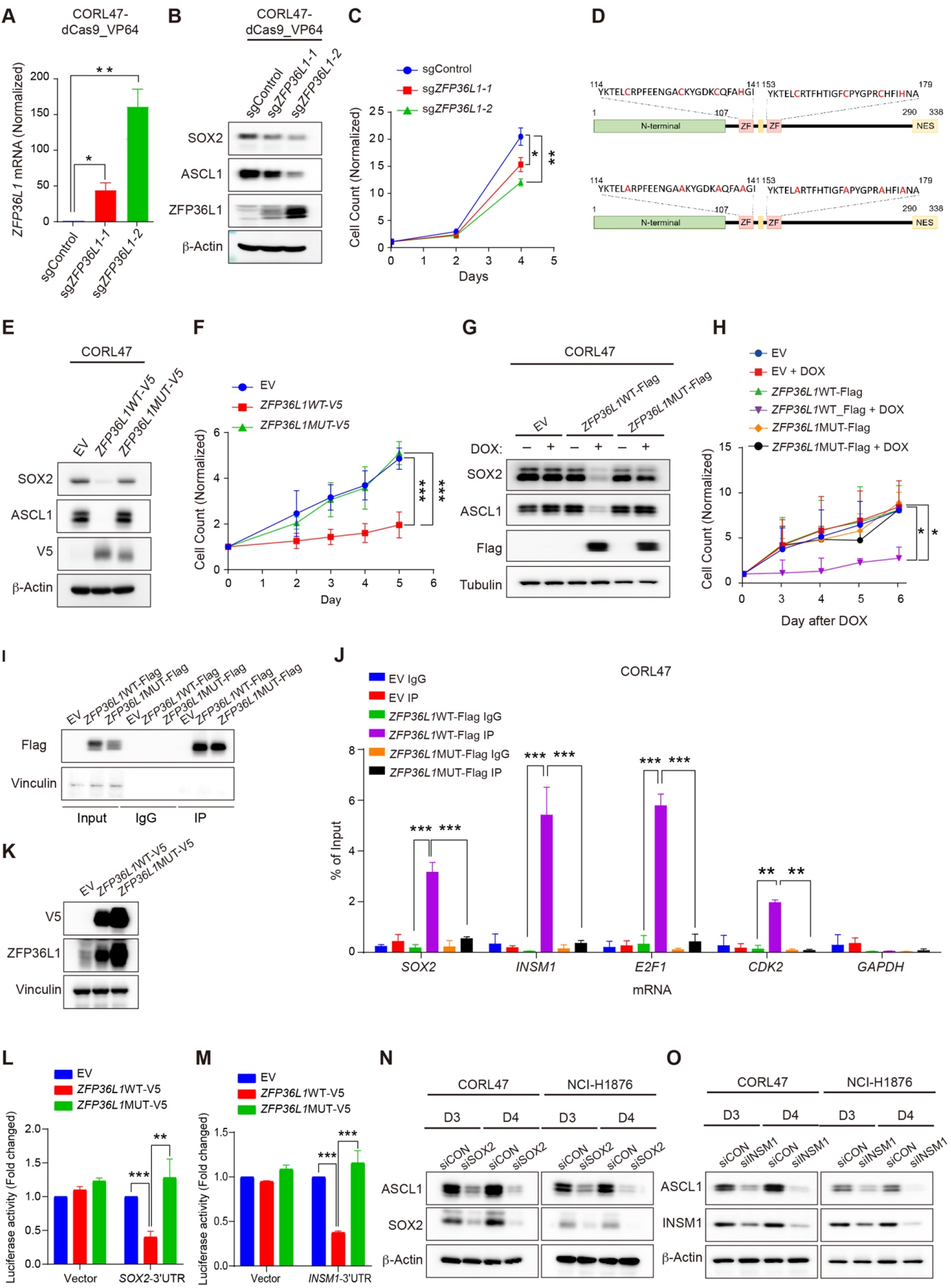
ZFP36L1 Blocks Neuroendocrine Differentiation and Proliferation in SCLC in part Through its Ability to Bind and Degrade the 3’ UTRs of SOX2 and INSM1. (**A and B**) RT-qPCR and immunoblot analysis of CORL47 dCas9-VP64 cells that were infected with sgRNAs targeting the promoter of ZFP36L1 (sgZFP36L1-1 or sgZFP36L1-2) or a non-targeting sgRNA (sgControl). n=3 biological replicates. (**C**) Proliferation assays of the cells in A. n=3 biological replicates. (**D**) Schematic showing the cysteine, C, sites in the Zinc-finger RNA-binding domains of ZFP36L1 that were mutated to alanines, A, to disrupt the RNA-binding activity of ZFP36L1. (**E**) Immunoblot analysis of CORL47 cells stably infected with ZFP36L1WT- V5, the ZFP36L1 mRNA-binding mutant-V5 (ZFP36L1MUT-V5), or the corresponding empty vector (EV). (**F**) Proliferation assays of the cells in E. n=3 biological replicates. (**G**) Immunoblot analysis of CORL47 cells stably infected with a doxycycline inducible (DOX-ON) lentivirus encoding ZFP36L1WT-FLAG, ZFP36L1MUT-FLAG, or the corresponding EV. Cells were treated with DOX (1 μg/mL) for 3 days where indicated. (**H**) Proliferation assays of the cells in G grown in the presence or absence of DOX (1 μg/mL) for the days indicated. n=3 biological replicates. (**I**) Immunoblot analysis after immunoprecipitation (IP) of ZFP36L1WT-FLAG, ZFP36L1MUT-FLAG, or EV CORL47 cells treated with DOX (1 μg/mL) for 3 days relative to input. (**J**) RNA quantitation relative to input after RT-qPCR from IP experiments in I with primers specific to SOX2, INSM1, E2F1, CDK2 and GAPDH. n=4 biological replicates. (**K**) Immunoblot analysis of 293T cells stably infected with ZFP36L1WT-V5, ZFP36L1MUT-V5, or the EV. (**L and M**) 3’ UTR luciferase reporter assay in 293T cells from K that were transfected with a SOX2- 3’UTR luciferase reporter (**L**), a INSM1-3’UTR luciferase reporter (**M**), or the corresponding EV (vector), and bioluminescence (BLI) was measured 24 hours later. The BLI signal of each condition is first normalized to the cell count and then normalized to the plx304 EV of each reporter. n=3 biological replicates. (**N and O**) Immunoblot analysis of CORL47 and NCI-H1876 cells transfected with siGenome SMARTPool targeting SOX2 (siSOX2) or non-targeting siRNA control (siCON) (**N**) or the siGenome SMARTPool targeting INSM1 (siINSM1) or siCON (**O**) at the times indicated after transfection. n=3 biological replicates. For all panels, *=p<0.05, **=p<0.01, ***=p<0.001.

### ZFP36L1 Directly Binds SOX2 and INSM1 to Block Neuroendocrine Differentiation

We next sought to identify candidate ZFP36L1 mRNA substrates that control SCLC neuroendocrine differentiation and cellular proliferation. Analysis of a select set of neuroendocrine transcription factors showed that the 3’ UTRs of SOX2 and INSM1, but not of ASCL1, contain the canonical AUUUA binding motifs for ZFP36L1 (Extended Data Fig. S9A). SOX2 and INSM1 are dependencies in some SCLC cell lines^34^. We therefore hypothesized that ZFP36L1 binding and downregulating SOX2 and INSM1 mRNAs could be responsible for ZFP36L1’s ability to block neuroendocrine differentiation and proliferation. To test this, we performed RNA-binding protein immunoprecipitation (RIP) for FLAG-tagged ZFP36L1 WT or the ZFP36L1-mRNA binding mutant in CORL47 cells and performed RT-qPCR for SOX2 and INSM1 along with previously identified ZFP36L1 targets including E2F1 and CDK2^33^. ZFP36L1 WT, but not the ZFP36L1-mRNA binding mutant, bound to the mRNAs of SOX2, INSM1, E2F1, and CDK2 (Fig. 5I,J). These results were confirmed using constitutively expressed V5-tagged ZFP36L1 (Extended Data Fig. S9B-C). We next asked whether ZFP36L1 WT directly binds and downregulates the 3’ UTRs of SOX2 and INSM1. To do this, we first engineered 293T cells to exogenously express ZFP36L1 WT or the ZFP36L1 mRNA-binding mutant (Fig. 5K) and transfected these cells with the 3’ UTR of SOX2 or INSM1 fused to RenSP luciferase or an empty vector control. ZFP36L1 WT expression, but not expression of the ZFP36L1 mRNA-binding mutant, dramatically reduced the bioluminescence signal of the SOX2 or INSM1 3’ UTR with no effect on the empty vector control (Fig. 5L,M). Treatment of CORL47 and NCI-H1876 SCLC cell lines with SOX2 siRNAs or INSM1 siRNAs potently decreased ASCL1 protein levels compared to the non-targeting siRNA control (Fig. 5N,O) showing that SOX2 or INSM1 loss phenocopies ZFP36L1 re-expression with respect to ASCL1. Together, these data demonstrate that ZFP36L1 directly binds and degrades the 3’ UTRs of SOX2 and INSM1.

### Restoring ZFP36L1 Expression Induces a Non-Neuroendocrine Phenotype with “Inflammatory” Features

An “inflammatory” subtype of SCLC that lacks neuroendocrine markers, restores MHC class I antigen presentation, and is more responsive to immune checkpoint blockade has recently been described^35–37^. Since ZFP36L1 and ASCL1 are expressed in mutually exclusive subpopulations in mouse SCLC tumors (Fig. 3G) and ZFP36L1 re-expression causes loss of neuroendocrine differentiation in SCLC cell lines (Fig. 5), we hypothesized that cells with restored ZFP36L1 expression would acquire markers that resemble the “inflammatory” subtype. To test this, we used CORL47 expressing 2 independent sgRNAs that activate endogenous ZFP36L1 or a non-targeting sgRNA as a control (see Fig. 5A) and performed RNA-sequencing followed by GSEA. Differential expression analysis demonstrated that endogenous ZFP36L1 activated cells highly expressed canonical HLA genes including HLA-B and HLA-C (Fig. 6A, B). Consistent with this, GSEA showed striking enrichment of gene sets of epithelial-mesenchymal transition and interferon gamma response in ZFP36L1 activated cells (Fig. 6C, D). Similar to our findings above (see Fig. 5), ZFP36L1 activation caused loss of neuroendocrine markers and ASCL1 target genes (Fig. 6E, F). Furthermore, RNA-sequencing data from human SCLCs with restored MHC class I antigen presentation^37^ had significantly higher ZFP36L1 expression compared to human SCLCs with silenced MHC class I (Fig. 6G). These results demonstrate that restoring ZFP36L1 can functionally induce a non-neuroendocrine state that resembles the “inflammatory” subtype of SCLC.

**Fig. 6:**
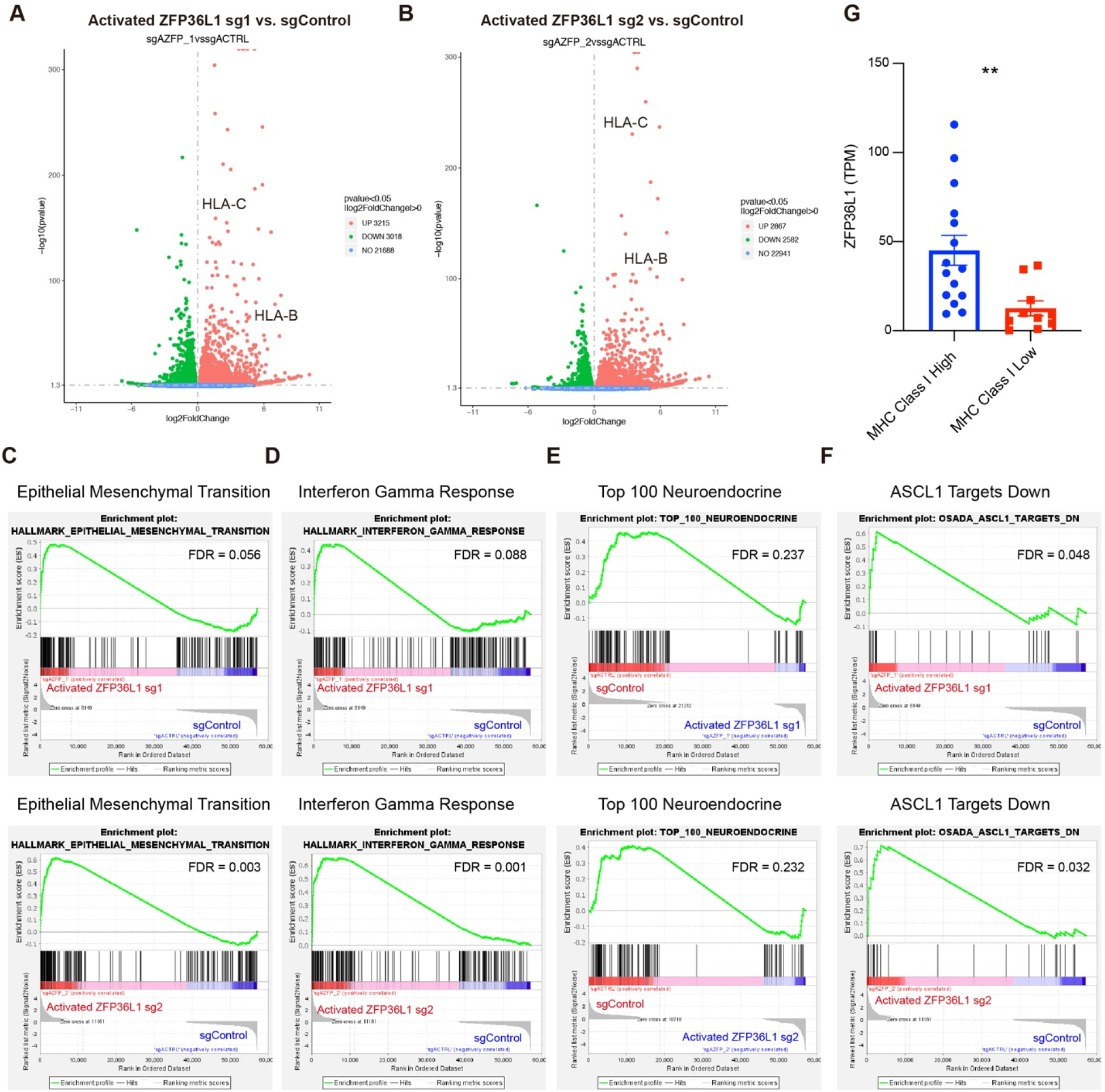
Restoring Endogenous ZFP36L1 Expression Promotes Non-Neuroendocrine Plasticity Toward an “Inflammatory” Phenotype. (**A and B**) Volcano plot of RNA-sequencing (RNA-seq) data showing the log2 fold change and - Log10 p-value of gene expression comparing CORL47 dCas9-VP64 cells expressing sgRNAs targeting the promoter of ZFP36L1 [sgZFP36L1-1 (**A**) or sgZFP36L1-2 (**B**)] to activate endogenous ZFP36L1 expression to a non-targeting sgRNA (sgControl). n=2 biological replicates. (**C-F**) GSEA of RNA-seq data in A (**Top**) or B (**Bottom**) of Epithelial-Mesenchymal Transition (**C**), Interferon Gamma Response (**D**), Top 100 Neuroendocrine Genes (**E**), and ASCL1 Target Genes Down (**F**). FDR q-values are indicated. (**G**) Quantification of RNA-sequencing data from human SCLC tumors with confirmed high or low MHC class I surface protein expression by IHC^37^. **=p<0.01.

### ZFP36L1 Induction Correlates with Sensitivity to ORY-1001 in SCLC Patient-Derived Xenograft Models

We observed a positive correlation between ZFP36L1 induction and inhibition of cellular proliferation after LSD1 inhibitor treatment (see Fig. 2) suggesting that ZFP36L1 induction could predict response to LSD1 inhibitors. To further investigate this, we interrogated a publicly available RNA-sequencing dataset of SCLC patient-derived xenograft (PDX) models that were treated with ORY-1001 and displayed a variable response to ORY-1001^7^. In these data, ORY- 1001 induced a near complete response in the FHSC04 model, while most other models were partially sensitive, and the LX33 model was completely resistant^7^. We first included the hits from each arm of our ORY-1001/KDM5-C70 combination screen with a q-value cut-off of less than 0.25 as a gene set and asked whether there were differences in enrichment of the hits measured using an area under the curve (AUC) algorithm (see Methods) in the ORY-1001 sensitive vs. resistant PDX models at baseline. We found enrichment of expression of screen hits in the FHSC04 highly sensitive model and relative depletion of screen hits in the LX33 resistant model (Fig. 7A and Extended Data Fig. S10B-D) with no changes in AUC in the enrichment hits from the untreated DMSO arm (Extended Data Fig. S10A) suggesting that high baseline expression of genes that are required for ORY-1001 sensitivity positively correlates with the baseline sensitivity of SCLC PDX models to ORY-1001. We next used the RNA-sequencing data set from these SCLC PDX models treated with ORY-1001 or DMSO *ex vivo* and performed an unsupervised hierarchical clustering analysis of mRNAs induced by ORY- 1001 relative to DMSO for each model. Strikingly, we found that *ZFP36L1* and *REST* are highly upregulated upon ORY-1001 treatment across all models, but that the magnitude of ZFP36L1 induction correlates strongly with baseline ORY-1001 sensitivity of each model with the highest magnitude of ZFP36L1 induction in the FHSC04 highly sensitive model and no ZFP36L1 induction in the LX33 resistant model (Fig. 7B and Extended Data Fig. S11A-C). In contrast, the LSD1 target gene REST^22^ was induced across all models and its magnitude of induction did not correlate with ORY-1001 sensitivity (Fig. 7B and Extended Data Fig. S11A-C). Furthermore, ZFP36L1 mRNA expression was one of the most significantly upregulated genes upon ORY- 1001 treatment *in vivo* in the FHSC04 highly sensitive model (Extended Data Fig. S11D,E). Together, these data suggest that ZFP36L1 induction after ORY-1001 treatment is a candidate biomarker to predict response to ORY-1001.

**Fig. 7:**
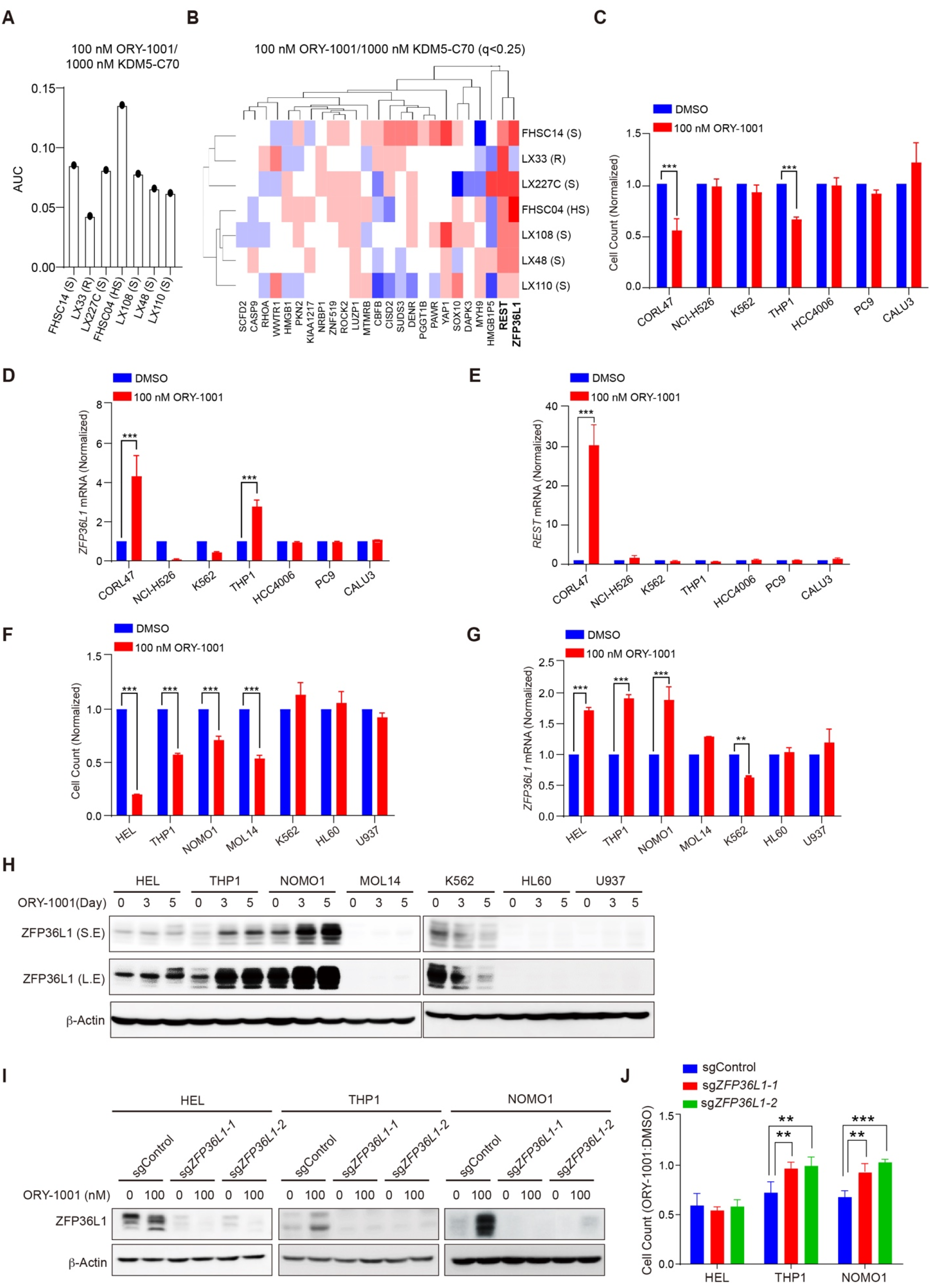
ZFP36L1 Induction after LSD1 Inhibitor Treatment Correlates with LSD1 Inhibitor Sensitivity in SCLC and AML. (**A**)Area under the curve (AUC) enrichment analysis of hits from our ORY-1001/KDM5-C70 resistance screen (in Fig. 1F) of RNA-seq data from untreated patient-derived xenograft (PDX) models of SCLC. (**B**) Unsupervised hierarchical clustering of hits from our ORY-1001/KDM5- C70 resistance screen (in Fig. 1F) of RNA-seq data from SCLC PDX models treated ex-vivo with ORY-1001 or DMSO. Red denotes genes with high expression, and blue denotes genes with low expression. (**C-D**) Quantitation of cell counts (**C**) and RT-qPCR for ZFP36L1 (**D**) or REST (**E**) of CORL47 (SCLC), NCI-H526 (SCLC), K562 (Leukemia), THP1 (Leukemia), PC9 (NSCLC), HCC4006 (NSCLC) and CALU3 (NSCLC) cells treated with ORY-1001 (100 nM) or DMSO for 6 days. For C-E, n=3 biological replicates. (**F-H**) Cell proliferation (**F**), RT-qPCR (**G**), and immunoblot analysis (**H**) of the leukemia cell lines indicated treated with ORY-1001 (100 nM) compared to cells treated with DMSO. For F and G, cells were treated with ORY-1001 for 3 days. n=3 biological replicates. (**I and J**) Immunoblot analysis (**I**) and quantitation of cell counts (**J**) of HEL, THP1, or NOMO1 cells first infected with lentivirus that expressed Cas9 and a sgRNA targeting ZFP36L1 (sgZFP36L1-1 or sgZFP36L1-2) or a non-targeting sgRNA (sgControl) and then treated with ORY-1001 (100 nM) or DMSO for 3 days. Data are plotted as the cell counts of ORY-1001-treated samples relative to DMSO-treated samples. n=3 biological replicates. For all panels, *=p<0.05, **=p<0.01, ***=p<0.001.

### ZFP36L1 is Induced by ORY-1001 and is Required for ORY-1001 Sensitivity in AML Cell Lines

We then asked whether ZFP36L1 induction correlates more broadly with LSD1 inhibitor sensitivity across other cancer cell lines. To test this, we first treated SCLC, NSCLC, and AML cell lines with ORY-1001 and performed RT-qPCR for ZFP36L1 and REST and measured cellular proliferation. In our initial screening panel, ORY-1001 inhibited proliferation in only 2 cell lines: the CORL47 SCLC cell line and the THP1 AML cell line (Fig. 7C). Strikingly, these cell lines were the only ones where ZFP36L1 mRNA was induced (Fig. 7D). In contrast, REST mRNA was induced in CORL47 cells but not in THP1 cells (Fig. 7E), suggesting that REST upregulation tracks with the SCLC lineage, but not with ORY-1001 sensitivity across cancer cell lines. To further explore whether ZFP36L1 induction correlates with ORY-1001 sensitivity in AML, we examined additional AML cell lines and found that 4 of the 7 AML cell lines were sensitive to ORY-1001 (Fig. 7F). ZFP36L1 mRNA and protein were induced in 3 (HEL, THP1, and NOMO1) of the 4 ORY-1001 sensitive cell lines (Fig. 7G,H). ORY-1001 treatment did not induce ZFP36L1 in the 3 resistant AML cell lines (K562, HL60, U937) (Fig. 7G,H). Similar to our findings in SCLC (see Fig. 4), CRISPR-mediated ZFP36L1 inactivation caused near complete resistance to ORY-1001 in 2 of the 3 AML cell lines (Fig. 7I,J) with the exception being HEL cells which only showed weak induction of ZFP36L1 upon ORY-1001 treatment (Fig. 7G,H). Overall, our data demonstrates that ZFP36L1 induction highly correlates with ORY-1001 sensitivity and that ZFP36L1 is functionally required for ORY-1001 sensitivity in SCLC and AML.

## Discussion

SCLC and AML are selectively sensitive to LSD1 inhibition^7–10^ with ongoing clinical trials testing LSD1 inhibitors in SCLC and AML patients^11^. Previous work demonstrated that LSD1 and KDM5 inhibitors promote loss of ASCL1 and neuroendocrine differentiation through a NOTCH-dependent mechanism^7, 15^. Consistent with previous findings, our CRISPR/Cas9 LSD1+KDM5 inhibitor resistance screen found that the NOTCH-target gene REST, which is necessary and sufficient for non-neuroendocrine plasticity in SCLC^23^ and an LSD1 target gene in SCLC^7, 22^, is required for LSD1+KDM5 inhibitor sensitivity. This suggests that LSD1+KDM5’s NOTCH-dependent control of ASCL1 and neuroendocrine differentiation converges on its ability to repress REST. Our screen discovered several genes required for LSD1 inhibitor sensitivity and SCLC neuroendocrine differentiation that were not previously linked to LSD1 in SCLC. For example, two top hits from our screen were *YAP1* and *WWTR1*, which we found are de- repressed upon LSD1 inhibition. YAP1 is expressed in a fraction of non-neuroendocrine SCLCs that lack ASCL1 expression^3, 38^ and repressed across several neuroendocrine cancers to sustain their neuroendocrine phenotype^39^. Our findings suggest that restoring YAP1 expression through LSD1 inhibition contributes to the non-neuroendocrine phenotype that emerges after LSD1 inhibition and suggests that LSD1 inhibition could induce the conversion of ASCL1-positive SCLCs to a YAP1-positive state.

We primarily focused on the mRNA-binding protein ZFP36L1 since ZFP36L1 was a top hit in all 4 arms of our screen, is selectively repressed in SCLC, and, to our knowledge, has never been studied in SCLC. We found that LSD1 binds and represses ZFP36L1 expression. Upon LSD1 inhibition, ZFP36L1 expression is restored, which blocks neuroendocrine differentiation and SCLC proliferation. ZFP36L1 has previously been shown to function as a tumor suppressor in other cancers through its ability to bind AU-rich elements in the 3’ UTR of mRNAs leading to mRNA degradation and tumor suppression^29, 33^. Previous identified mRNA targets of ZFP36L1 include mRNAs that drive proliferation in other cancers including Cyclin D, E2F1, and NOTCH^29, 33^. Our results are consistent with these studies identifying ZFP36L1 as a tumor suppressor in SCLC, where it also possesses the ability to bind and downregulate mRNAs that promote SCLC neuroendocrine differentiation including INSM1 and SOX2. SOX2 is required for SCLC proliferation ^6^ and is necessary for neuroendocrine lineage plasticity upon resistance to anti-androgen directed therapy in prostate cancer^32^. INSM1 is a neuroendocrine transcription factor marked by superenhancers^40^, is a dependency in some SCLC cell lines^34^, and is used as an immunohistochemical marker to diagnose SCLC^41^. Together with these previous findings, our data nominate SOX2 and INSM1 mRNAs as critical targets of ZFP36L1 required for its ability to block neuroendocrine differentiation and SCLC proliferation. ZFP36L1’s control of SCLC neuroendocrine is likely not limited to its regulation of SOX2 and INSM1. Future studies will focus on comprehensively identifying ZFP36L1 target mRNAs that control SCLC neuroendocrine differentiation and survival.

ZFP36L1 is highly repressed in SCLC, but is expressed in most other solid tumors suggesting that ZFP36L1 repression is selected for during SCLC tumorigenesis. ZFP36L1 is likewise repressed in other neuroendocrine tumors including neuroblastoma, that like SCLC, are dependent on ASCL1^30^, suggesting that ZFP36L1 repression is a common feature across high-grade neuroendocrine tumors. We hypothesize that ZFP36L1 repression is required to sustain SCLC neuroendocrine differentiation during SCLC tumorigenesis. Consistent with this, we identified subpopulations in mouse SCLC tumors that express ZFP36L1 and have relatively lower ASCL1 expression compared to the bulk of cells within SCLC tumors that are ZFP36L1 negative and highly express ASCL1. We also found that restoring endogenous ZFP36L1 expression caused loss of neuroendocrine differentiation and promoted a non-neuroendocrine “inflammatory” phenotype with high expression of HLA genes and other inflammatory markers and that ZFP36L1 expression is enriched in the small fraction of human SCLCs that restore MHC class I antigen presentation. Recently, an “inflammatory” subtype of human SCLC has been identified, which has increased immunogenicity and is more responsive to immune checkpoint blockade^35–37^. Our results suggest that restoring ZFP36L1 expression in SCLC can drive plasticity toward this “inflammatory” subtype, which could be leveraged therapeutically. We found that LSD1 inhibition is one mechanism by which ZFP36L1 expression can be restored. Given the potent growth suppressive effects we observed after ZFP36L1 re-expression and ability to promote an inflammatory-like state, it will be important to determine whether inhibition of other repressive epigenetic modifiers such as EZH2 and KDM5A that have been previously linked to the SCLC neuroendocrine phenotype, SCLC survival, and immunogenicity^15, 37, 42–44^ also restore ZFP36L1 expression and if so, whether are additive/syngergistic with LSD1 inhibition. This could lead to combination therapeutic strategies that cooperatively restore ZFP36L1 promoting tumor suppression and increasing tumor immunogenicity.

ASCL1 is required for SCLC tumorigenesis and survival^13, 14^. Recent evidence demonstrates that MYC can drive SCLC plasticity from an ASCL1 to a NEUROD1 state and eventually into a non-neuroendocrine phenotype that lacks both ASCL1 and NEUROD1^45^. MYC can also drive SOX9-driven tumors when ASCL1 is genetically-inactivated^46^ suggesting that SCLCs can promote plasticity through MYC and become ASCL1-independent. We have identified a new mechanism of plasticity in SCLC through ZFP36L1-dependent regulation of mRNA stability that promotes non-neuroendocrine plasticity with increased HLA expression and is required for the acute sensitivity to LSD1 inhibitors in SCLC cell lines. It is unknown whether non-neuroendocrine cells with restored ZFP36L1 will emerge as a delayed resistance mechanism to evade their neuroendocrine dependence. It will be important to determine whether resistant human SCLCs from patients enrolled in LSD1 inhibitor clinical trials have a neuroendocrine or non-neuroendocrine inflammatory phenotype.

Both SCLC and AML are selectively sensitive to LSD1 inhibitors and have repressed ZFP36L1 expression at baseline. We found that LSD1 inhibitors in both SCLC and AML induce ZFP36L1 expression and require ZFP36L1 for their anti-proliferative effects. A previous study demonstrated a convergence between AML and SCLC with respect to therapeutic vulnerabilities^47^ and our data on the LSD1-ZFP36L1 pathway in SCLC and AML is another example of their convergence. Interestingly, recent data from an unbiased CRISPR/Cas9 screen to identify genes that promote differentiation in AML found that LSD1 and the ZFP36L1 paralog, ZFP36L2, were required to sustain AML dedifferentiation^48^. Analogous to our findings showing that ZFP36L1 promotes non-neuroendocrine differentiation in SCLC, they found that AMLs induce ZFP36L1 to differentiate. Furthermore, they found that ZFP36L2 is an AML dependency and degrades the mRNAs of ZFP36L1 and its paralog ZFP36, to block AML differentiation. Our study did not explore the ZFP36L1 paralogs ZFP36 and ZFP36L2 since ZFP36L1 was the only paralog that scored in our screen. The interplay and functional roles of ZFP36L1 paralogs on SCLC neuroendocrine differentiation will be explored in future studies.

LSD1 inhibitors show activity in preclinical models of SCLC and AML and clinical trials are ongoing. However, even in preclinical models, there is differential sensitivity among models suggesting the need for a predictive biomarker for patient selection. We found that a limited gene set of hits from our CRISPR/Cas9 screen was enriched in the highly sensitive SCLC PDX model and relatively depleted in the resistant model. However, this was a limited data set and larger number of PDX models, or preferably patient samples, are needed. We were unable to find a single biomarker that was predictive of response. Our data confirms previous findings that ASCL1 down-regulation after LSD1 inhibitor treatment correlates with sensitivity^7^, but ASCL1 expression alone was not predictive of LSD1 inhibitor sensitivity. We found that ZFP36L1 induction after LSD1 inhibitor treatment highly correlated with response in SCLC PDX models and SCLC cell lines. Future studies should focus on identifying predictive biomarkers to identify subsets of SCLC patients that are more likely to have benefit from LSD1 inhibitors.

## Acknowledgements

We thank the members of the Oser, Barbie, and Janne laboratories for helpful discussions, and Leslie Duplaquet for critical reading of the manuscript.

## Funding

M.G.O. is supported as a William Raveis Charitable Fund Clinical Investigator of the Damon Runyon Cancer Research Foundation (CI-101-19), by an NCI/NIH K08 grant (no. K08CA222657), and the Kaplan Family Fund.

## Authors’ Contributions

Conceptualization: H.Y.C and M.G.O.; Methodology: H.Y.C, Y.T.D., and M.G.O.; Investigation: H.Y.C, Y.T.D., Y.L., E.W., T.D. and M.G.O. Formal Analysis: H.Y.C, Y.T.D., Y.L., A.V., E.W., T.D. and M.G.O. Writing: H.Y.C. and M.G.O.; Review and Editing: All authors; Funding Acquisition: M.G.O.

## Competing Interests

M.G.O. has sponsored research agreements (SRAs) with Takeda, Eli Lilly, Novartis, and Bristol Myers Squibb. None of these SRAs were used to fund this work.

## Data and Materials Availability

Raw Data and Log normalized data from the CRISPR screen is included as Extended Data Table 1. Data from RNA sequencing experiments will be deposited in the GEO database prior to publication. All other data and materials can be requested from the corresponding author upon reasonable request.

## Methods

### Cell Lines and Cell Culture

NCI-H1876 (obtained 11/2016), NCI-H1092 (obtained in 11/2018), NCI-H2081 (obtained in 11/2018), NCI-H526 (obtained 10/2019), NCI-H1618 (obtained 1/2020), NCI-H2066 (obtained 1/2020) and 293FT cells were originally obtained from American Type Culture Collection (ATCC). CORL47 were obtained from Sigma (11/2018). NCI-H69 cells were a kind gift from Dr. Kwok-kin Wong’s laboratory (New York University, obtained 8/2014). HEL, THP1 NOMO1, MOL14, K562, HL60 and U937 were a kind gift form Dr. Julie-Aurore Losman’s laboratory at DFCI (10/2020). HCC4006, CALU3, and PC9 cells were a kind gift Dr. Pasi Janne’s laboratory at DFCI (7/2020).

CORL47, NCI-H526, NCI-H69, HCC4006, CALU3, PC9, HEL, THP1, NOMO1, MOL14, K562, HL60, and U937 cells were maintained in RPMI-1640 media supplemented with 10 % fetal bovine serum (FBS), 100 U/mL penicillin (P), and 100 μg/mL streptomycin (S). 293FT cells were maintained in DMEM media with 10% FBS and P/S. NCI-H1092, NCI-H1618, NCI-2066, NCI-H1876, and NCI-H2081 were maintained in DMEM/F12 with 5% FBS, P/S, and HITES [10 nM hydrocortisone, Insulin-Transferrin-Selenium (Gemini), and 10 nM beta-estradiol].

Early passage cells of all the cell lines listed above were frozen using Bambanker’s freezing media (Bulldog Bio) and were maintained in culture no more than 4 months where early passage vials were thawed. Where indicated, the following chemicals (stored at -20°C or -80°C) were also added to the media: doxycycline (stock 1 mg/mL in H_2_0), KDM5-C70 (Xcessbio #M60192-2, stock 10 mM in DMSO), or ORY-1001 (Selleck #S7795, stock 10 mM in DMSO), or CC-90011 (Selleck #S0356, stock 10 mM in DMSO).

### sgRNA Cloning

sgRNA sequences were designed using the Broad Institutes sgRNA designer tool (http://portals.broadinstitute.org/gpp/public/analysis-tools/sgrna-design) and synthesized by IDT technologies. The sense and antisense oligonucleotides were mixed at equimolar ratios (0.25 nanomoles of each sense and antisense oligonucleotide) and annealed by heating to 100°C in annealing buffer (1X annealing buffer 100 mM NaCl, 10 mM Tris-HCl, pH 7.4) followed by slow cooling to 30°C for 3 hours. The annealed oligonucleotides were then diluted at 1:400 in 0.5X annealing buffer.

For CRISPR/Cas9 loss of function experiments, the annealed oligos were ligated into plentiGuide-sgRNA-P2A-mCherry-Puro^50^ for all experiments in SCLC and AML cell lines or lentiCRISPR V2-Neo (Addgene #52961 where the puromycin resistance gene was replaced with the G418 resistance gene) for experiments in PC9 cells. For CRISPR activation experiments, the annealed oligos were ligated into pXPR_502 (Addgene #96923) vectors. Ligations were done with T4 DNA ligase for 2 hours at 25°C. The ligation mixture was transformed into HB101 competent cells. Ampicillin-resistant colonies were screened by restriction digestion of miniprep DNAs and subsequently validated by DNA sequencing.

The following sgRNA oligos were used for plentiGuide-sgRNA-P2A-mCherry-Puro vector for CRISPR knockout experiments:

human ZFP36L1-1 sense (5′-CACCGGCTGTCTCGCGAGCTCAGAG-3′),

human ZFP36L1-1 anti-sense (5′-AAACCTCTGAGCTCGCGAGACAGCC-3′),

human ZFP36L1-2 sense (5′-CACCGAGGGGCAAAAGCCGATGGTG-3′),

human ZFP36L1-2 anti-sense (5′-AAACCACCATCGGCTTTTGCCCCTC-3′),

human ZFP36L1-3 sense (5′-CACCGAAACGGTGCCTGTAAGTACG-3′),

human ZFP36L1-3 anti-sense (5′-AAACCGTACTTACAGGCACCGTTTC-3′),

human ZFP36L1-4 sense (5′-CACCGGGGTGACTGAGTGCCTCCGA-3′),

human ZFP36L1-4 anti-sense (5′-AAACTCGGAGGCACTCAGTCACCCC-3′),

human ZFP36L1-5 sense (5′-CACCGAGTTGAGCATCTTGTTACCC-3′),

human ZFP36L1-5 anti-sense (5′-AAACGGGTAACAAGATGCTCAACTC-3′),

human ZFP36L1-6 sense (5′-CACCGTGCCGCACCTTCCACACCAT-3′),

human ZFP36L1-6 anti-sense (5′-AAACATGGTGTGGAAGGTGCGGCAC-3′)

human NO_SITE_110 sense (5′-CACCGCTAATCACGACCTCACCCTA-3′),

human NO_SITE_110 anti-sense (5′-AAACTAGGGTGAGGTCGTGATTAGC-3′),

The following sgRNA oligos were used for pXPR_502 vector for CRISPR activation experiments:

human ZFP36L1-1 sense (5′-CACCGCTGGAGTTGAATCAGCCATG-3′),

human ZFP36L1-1 anti-sense (5′-AAACCATGGCTGATTCAACTCCAGC -3′),

human ZFP36L1-2 sense (5′-CACCGATCAGCCATGCGGTCAGGCG -3′),

human ZFP36L1-2 anti-sense (5′-AAACCGCCTGACCGCATGGCTGATC -3′),

human non-targeting_C0111 sense (5′-CACCGGGAGGCTAAGCGTCGCAA -3′),

### Lentiviral cDNA and 3’UTR Expression Vector Cloning

To make the DOX-On pTripZ *ZFP36L1-WT* cDNA and pLX304-CMV ZFP36L1-WT cDNA, pcDNA3.1 mcherry-ZFP36L1 (Addgene #121140) was used as a template for overhang PCR that introduced attB1 and attB2 sites onto the 5’ and 3’ ends of ZFP36L1 (sense primer 5’- GGGGACAAGTTTGTACAAAAAAGCAGGCTTCGCCACCATGACCACCACCCTCGTGTCTGCC and anti-sense primer 5’- GGGGACCACTTTGTACAAGAAAGCTGGGTCGTCATCTGAGATGGAAAGTCTG-3’). The PCR product was then gel extracted per the manufacturer’s instructions (Qiagen). To make the DOX-On pTripZ *ZFP36L1-MUT* cDNA and pLX304-CMV ZFP36L1-MUT cDNA, the ZFP36L1 mRNA-binding mutant (referred to hereafter as ZFP36L1-MUT) was made by mutating 8 cysteines or histidines to alanines to disrupt the mRNA binding activity^33^, designed *in silico* using twist.com to introduce the intended cysteine (C) or histidine (H) to alanine (A) non-synonymous mutations and introducing synonymous mutations as necessary to achieve synthesis. AttB1 and attB2 sites onto the 5’ and 3’ ends, respectively, for gateway cloning. The ZFP36L1-MUT construct was then synthesized as a double-stranded DNA G-Block by IDT technologies.

Purified ZFP36L1-WT and ZFP36L1-MUT double-stranded DNA fragments containing attB1 and attB2 sites were introduced into the pDONR223 vector by homologous recombination using BP clonase (Life Technologies #11789020) for 1 hour at 25 °C according to the manufacturer’s instruction. The reaction mixture was then transformed at a ratio of 1:10 (reaction volume/volume competent cells) into HB101 competent cells (Promega). Spectinomycin-resistant colonies were screened by restriction digestion of miniprep DNA and subsequently validated by DNA sequencing. A homologous recombination reaction was then performed using LR Clonase II (Life Technologies #11791100) with the pDONR223-ZFP36L1- WT or pDONR223-ZFP36L1-MUT plasmids and DOX-On pTripZ-NEO DEST or pLX304-CMV DEST vectors that were both modified to be used as destination vectors for gateway recombination cloning. The LR reaction was transformed into HB101 competent cells (Promega). Kanamycin-resistant (pTripZ-NEO) or ampicillin-resistant (PLX304-CMV) colonies were screened by restriction digestion of miniprep DNA and subsequently validated by DNA sequencing.

For pLightSwitch-h*INSM1*-3’UTR construction, genomic DNA was isolated from CORL47 cells using the QIAamp DNA Blood Mini Kit (Qiagen #51106) according to the manufacturer’s instructions. Nested PCR was performed by two successive PCR reactions using KOD Xtreme polymerase (EMD Millipore #71975) and following set of primers to amplify the genomic region of *INSM1-3’UTR*:

outer forward *INSM1-3’UTR* (5’-CAGGTGTTCCCCTGCAAGTA-3’),

outer reverse *INSM1-3’UTR* (5’-CCTGGTGGTTCCCTTTTTCC-3’),

inner forward *INSM1-3’UTR with NheI* (5’-GCTAGCAGCGCGCCCTCCACC-3’),

inner reverse *INSM1-3’UTR with XhoI*

(5’-CTCGAGGACTTTGAAAATATTTTATTTGAATAGTAAATAGTTATACAG-3’).

The final PCR product was then column purified using Qiagen’s gel extraction kit (Qiagen #28706X4). The pLightSwitch-h*INSM1*-3’UTR was then made by digesting the pLightSwitch-EV (SwitchGear Genomics) and the INSM1-3’UTR PCR product with NheI and XhoI for 2 h at 55°C and gel-band purifying the resulting linearized fragments. Ligations were performed with T4 DNA ligase overnight at room temperature. The ligation reaction was transformed into HB101 competent cells (Promega). Ampicillin-resistant colonies were screened by restriction digestion of miniprep DNA and subsequently validated by DNA sequencing. SOX2-3’-UTR was purchased from SwitchGear Genomics (#S808404).

Lenti-Cas9Blast (#52962) and dCas9-VP64_Blast (#61425) were purchased from Addgene.

### Lentivirus Production

Lentiviruses were made by Lipofectamine 2000-based co-transfection of 293FT cells with the respective lentiviral expression vectors and the packaging plasmids psPAX2 (Addgene #12260) and pMD2.G (Addgene #12259) at a ratio of 4:3:1. Virus-containing supernatant was collected at 48 and 72 h after transfection, pooled together (15 mL total per 10-cm tissue culture dish), passed through a 0.45-µm filter, aliquoted, and frozen at −80°C until use.

### Lentiviral Infection

The SCLC and AML cells were counted using a Vi-Cell XR Cell Counter (Beckman Coulter) and resuspended in 1 mL lentivirus with 8 μg/mL polybrene at the following concentrations in individual wells of a 12 well plate: 1 × 10^6^ cells/mL for NCI-H69, HEL, NOMO1 and THP1 cells, or 2 X10^6^ cells/mL for NCI-H1876 and CORL47 cells. The plates were then centrifuged at 448 x g for 2 hours at 30° C. 16 hours later the virus was removed and cells were grown for 72 hours before being placed under drug selection. Cells were selected in puromycin (0.5 μg/mL for SCLC cell lines and 1 μg/mL for AML cell lines), blasticidin (5 μg/mL), or G418 (500 μg/mL for SCLC and 1000 μg/mL for AML) and maintained in media containing puromycin (0.5 μg/mL), blasticidin (2.5 μg/mL), or G418 (250 μg/mL for SCLC cells and 500 μg/mL for AML cells).

### Immunoblot Analysis

∼1 × 10^6^ cells were centrifuged at 252 x g for 5 minutes at 4°C and the media was removed by gentle aspiration. The cell pellet was then washed once in 1 mL of ice-cold PBS, transferred to a 1.5 mL Eppendorf tube, and centrifuged at 252 x g for 5 minutes at 4°C. The PBS was carefully aspirated and the cell pellet was lysed in EBC lysis buffer (50mM Tris Cl pH 7.5, 250 mM NaCl, 0.5% NP-40, 10% Glycerol) supplemented with a protease inhibitor cocktail (Complete, Roche Applied Science, #11836153001) and phosphatase inhibitors (PhosSTOP Sigma #04906837001). Soluble extracts were quantified using the Bradford Protein Assay. The protein sample was boiled in sample buffer (3X is 6.7% SDS, 33% Glycerol, 300 mM DTT, and Bromophenol Blue), resolved by 10% SDS-polyacrylamide gel electrophoresis (SDS-PAGE), transferred onto nitrocellulose membranes, blocked in 5% milk in Tris-Buffered Saline with 0.1% Tween 20 (TBS-T) for 1 hour, and probed with the indicated primary antibodies overnight at 4°C. Membranes were then washed 3 times in TBS-T, probed with the indicated horseradish peroxidase-conjugated (HRP) secondary antibodies for 2 hours at room temperature, and washed 3 times in TBS-T. Bound antibodies were detected with enhanced chemiluminescence (ECL) western blotting detection reagents [Immobilon (Thermo Fisher Scientific, #WBKLS0500) or Supersignal West Pico (Thermo Fisher Scientific, #PI34078)].

The primary antibodies used were: rabbit α-ZFP36L1 (Cell Signaling, hBRF1/2, #2119, used at 1:1000), mouse α-INSM1 (SANTA CRUZ, #SC377428, used at 1:1000), rabbit α-SOX2 (Cell Signaling, #3579, used at 1:1000), rabbit α-ASCL1 (Abcam, # ab211327, used at 1:1000) mouse α-Vinculin (Sigma, hVIN-1, # V9131, used at 1:10,000), mouse α-Tubulin (Sigma, B-5-1-2, # T5168, used at 1:5000), mouse α-FLAG (Sigma, clone M2, #F1804, used at 1:2000), rabbit α-V5 (Cell Signaling, #13202, used at 1:1000) and mouse α-®-actin (Sigma, clone AC-15, #A3854, used at 1:25,000). The HRP conjugated secondary antibodies were Goat α-Mouse (Jackson ImmunoResearch 115-035-003) and Goat α-Rabbit (Jackson ImmunoResearch 111-035-003) and were used at 1:5000.

### Positive-selection CRISPR/Cas9 ORY-1001 and ORY-1001/KDM5-C70 Resistance Screen

NCI-H1876 cells that had been infected with lenti-Cas9Blast (Addgene #52962) and subsequently maintained in Blasticidin were used for the screen. The screen was performed in 2 biological replicates. Cas9 expression was confirmed by immunoblot analysis and Cas9 activity was confirmed using a Cas9 GFP reporter [(pXPR_011 (Addgene #59702)] that showed near maximal editing 10 days after infection. On day 0, these NCI-H1876 Cas9Blast cells were counted. 1.5 × 10^8^ cells ( 500 cells/sgRNA anticipating an MOI of ∼0.3) were pelleted and resuspended at 2 × 10^6^ cells/mL in complete media containing 23 μl/mL of the whole genome Brunello sgRNA library (CP0041) lentivirus and 8 µg/mL polybrene, which yielded a multiplicity of infection (MOI) of ∼0.3, which was determined empirically in pilot experiments. The sgRNA Brunello whole genome library (CP0041) contains 77,441 sgRNAs targeting 4 sgRNAs per gene with 1000 non-targeting sgRNAs controls. The cells mixed with polybrene and lentivirus were then plated in 1 mL aliquots onto 12 well plates and centrifuged at 434 X g for 2 hours at 30°C. 16 hours later (day 1), the cells were collected, pooled, and centrifuged to remove the lentivirus and polybrene, and the cell pellet was resuspended in complete media at 1 × 10^6^ cells/mL and plated into 15 10 cm tissue culture treated plates. The cells were then cultured for 72 hours at which time (day 4) the media wash changed to media containing puromycin (0.5 µg/ml) to select for transduced cells. A parallel experiment was performed on day 4 to determine the MOI. To do this, the cells infected with the sgRNA library or mock-infected cells were plated at 1 × 10^6^ cells/mL in the presence or absence of puromycin. After 72 hours (day 7), cells were counted using the Vi-Cell XR Cell Counter and the MOI was calculated using the following equation: (# of puromycin-resistant cells infected with the sgRNA library/# total cells surviving without puromycin after infection with the sgRNA library) – (# of puromycin-resistant mock-infected cells/ # total mock-infected cells). The actual MOI was 0.42 for biological replicate 1 and 0.41 for biological replicate 2. On day 7 after MOI determination, the media was changed and replaced with fresh media again containing 0.5 µg/ml of puromycin.

On day 11, all puromycin-resistant cells were pooled, collected, and counted. A total of 4.4 X10^7^ cells for each condition (therefore maintaining at least 500 cells/guide) were plated at 1X10^6^ cells/mL into 2 15 cm tissue culture plates into complete media without puromycin that contained the following: 1. ORY-1001 (1 nM); 2. ORY-1001 (100 nM); 3. ORY-1001 (1 nM) + KDM5-C70 (1000 nM); 4. ORY-1001 (100 nM) + KDM5-C70 (1000 nM); 5. KDM5-C70 (1000 nM); or 6. DMSO. Note, a DMSO condition was only included for replicate 1. At the same time 4.4 × 10^7^ cells were collected from the pooled puromycin-resistant cells described above, washed in PBS, and the cell pellets were frozen for genomic DNA isolation for the initial timepoint prior drug selection. In parallel, mock infected cells were maintained at the same density for all drug conditions and carried in parallel for the entire screen described below. On day 14, the media was changed into fresh complete media containing fresh drug. On day 17, the cells were pooled, collected, counted, and replated at 4.4 X10^7^ cells for each condition containing fresh drug as was done on day 11. On days 20 and 23, the media was changed into fresh complete media containing fresh drug. On day 25, conditions 1,2,5, and 6 were trypsinized, counted, and resuspended at 1X10^6^ cells/mL (4.4 X10^7^ cells total) in complete media containing fresh drug into 2 15 cm plates. For conditions 3 and 4, the media was changed into fresh complete media containing fresh drug. On day 28, conditions 3 and 4 were trypsinized, counted, and all cells were resuspended in 44 mls of complete media containing fresh drug into 2 15 cm plates. For conditions 1,2,5, and 6, the media was changed into fresh complete media containing fresh drug. On day 31, the media was changed into fresh complete media containing fresh drug. On day 35, there was an obvious growth advantage for all conditions that contained ORY-1001 (1,2,3,4) of the cells infected with the sgRNA library and treated with drug compared to the mock infected cells treated with drug (see Fig. 1B) and therefore we ended the screen by collecting all remaining cells. Following completion of the screen, gDNA was isolated using a Qiagen Genomic DNA mini prep kit (cat. # 51104) for conditions 1,2,3, and 4 or with the Qiagen Genomic DNA maxi prep kit (cat. # 51192) for conditions 5 and 6 according to the manufacturer’s protocol with slight modifications made by the Broad Genetic Perturbation Platform. Raw Illumina reads were normalized between samples using: Log2[(sgRNA reads/total reads for sample) × 1e6) + 1]. The initial time point data (day 11) was then subtracted from the end time point after drug selection (day 35) to determine the relative enrichment of each individual sgRNA after the specified drug treatments. The subtracted data was then analyzed using Hypergeometric Analysis and the STARS algorithm on the GPP web portal (https://portals.broadinstitute.org/gpp/screener/) to determine the genes that were significantly enriched after ORY-1001 and/or KDM5-C70 treatment. A q-value cutoff of less than 0.25 calculated using the STARS analysis was used to call hits. The averaged data from 2 biological replicates were used for all analyses.

### Cell Proliferation Assays

For proliferation experiments with parental cell lines, cells were counted on day 0 using a Vi-Cell XR Cell Counter and plated in a tissue culture-treated 6-well plate at 100,000 cells/mL in 2 mL of complete media for CORL47 cells or at 250,000 cells/ml in 2 mLs of complete media for NCI-H1876 cells. AML cells were plated in tissue culture-treated 12 well plates at 100,000 cells/mL in 1 mL of media. For dose response assays with LSD1 inhibitors, the cells were treated with drug (ORY-1001 or CC-90011 at the concentrations indicated) for the number of days indicated.

For the proliferation experiments with the CORL47 ZFP36L1 CRISPR activation cells and all cells exogenously overexpressing ZFP36L1, cells were counted on day 0 using a Vi-Cell XR Cell Counter and plated in a 12-well at 200,000 cells/mL in 1 mL of media and counted on the days indicated. For the DOX-On inducible ZFP36L1-WT and ZFP36L1-MUT expression experiments, cells were grown in the presence or absence of Doxycycline (DOX) (1 μg/mL) as indicated after the initial cell counts were determined on day 0.

### Reverse-Transcriptase Quantitative PCR (RT-qPCR)

RNA was extracted using RNeasy mini kit (Qiagen #74106) according to the manufacturer’s instructions. RNA concentration was determined using the Nanodrop 8000 (Thermofisher Scientific). A cDNA library was synthesized using iScript Reverse Transcription Supermix for RT-qPCR (Biorad # 1708841) according to the manufacturer’s instructions. qPCR was performed using the LightCycler 480 (Roche) with the LightCycler 480 Probes Master Kit (Roche) and Taqman probes (ThermoFisher Scientific) according to the manufacturer’s instructions. The ÄÄC_T_ Method was used to analyze data. The C_T_ values for each probe were then normalized to the C_T_ value of *ACTB* for that sample. The data from each experiment was then normalized to the control to determine the relative fold change in mRNA expression. The following TaqMan probes were used: ZFP36L1 human (Hs00245183_m1), ASCL1 human (Hs04187546_g1), REST human (Hs05028212_s1), YAP1 human (Hs00902712_g1), WWTR1 human (Hs00210007_m1) and ACTB human (Hs01060665_m1).

### ChIP-qPCR Assays

CORL47 cells were plated at 1 × 10^6^ cells/mL in 10 mLs of complete media and NCI- H1876 cells were plated at 2.5 × 10^6^ cells/mL in 10 mLs of complete media containing ORY- 1001 (100 nM) or DMSO. The cells were harvested 6 days later. CORL47 cells were collected, and centrifuged. The supernatant was aspirated and the cells were washed once in PBS. For the NCI-H1876 cells, the media was aspirated and the cells were washed once in PBS. Both cell lines were then cross-linked by incubation with 2 mM disuccinimidyl glutarate (DSG) with rocking for 30 minutes at room temperature. The DSG was removed and the cells were fixed in 1% paraformaldehyde diluted in PBS for 10 minutes at 37°C. Excess formaldehyde was quenched by dropwise addition of freshly made Glycine at a final concentration 1.25 M followed by rocking the cells for an additional 5 minutes at room temperature. The paraformaldehyde/glycine was then removed and the cells were washed once with ice cold PBS. For CORL47 cells, the cells were centrifuged and washed once in ice-cold PBS. For NCI- H1876 cells, 5 mLs of ice-cold PBS was added and the cells were removed from the plate by scraping.

The cells were then centrifuged at 3000 x g for 5 minutes at 4° C and resuspended in 1 mL of lysis buffer (50 mM Hepes-NaOH pH 8, 140 mM NaCl, 1 mM EDTA, 10% glycerol, 0.5% NP-40, 0.25% TX-100 supplemented with a protease inhibitor cocktail and phosphatase inhibitors) and incubated by rotating for 10 minutes at 4°C. Intact nuclei were then collected by centrifugation in an Eppendorf microcentrifuge at 4500 rpm for 5 minutes at 4°C. The supernatant was then gently aspirated and the pellet was resuspended in 1 mL of wash buffer (10 mM Tris-HCl pH 8.0, 200 mM NaCl, 1 mM EDTA, supplemented with a protease inhibitor cocktail and phosphatase inhibitors) and incubated while rotating for 10 minutes at 4°C. The nuclei were again collected by centrifugation in an Eppendorf microcentrifuge at 4500 rpm for 5 minutes at 4°C. The supernatant was then aspirated and the tube was rinsed with 0.2 mL of shearing buffer (0.1% SDS, 1 mM EDTA, 10 mM Tris-HCl pH 8.0 supplemented with a protease inhibitor cocktail and phosphatase inhibitors). The nuclei were again collected by centrifugation in an Eppendorf microcentrifuge at 4500 rpm for 5 minutes at 4°C, the supernatant was carefully removed, and the pellet was resuspended in 1 mL shearing buffer and transferred to a 1 mL AFA tube (Covaris #520130).

The chromatin was then sonicated using a Covaris E220 sonicator at 100 peak incident power, 10% duty cycles, 200 cycles per burst, water level 15 (left indicator)/10 (right indicator) for 900 seconds. The samples were then transferred to 1.5 mL Eppendorf tubes and centrifuged in an Eppendorf microcentrifuge at 13,000 rpm for 5 minutes at 4°C. 0.9 mL of the supernatants were transferred to a new Eppendorf tube, to which TX-100 (final concentration of 1%) and NaCl (final concentration of 150 mM) were added.

20 μl of Dynabeads Protein G (ThermoFisher #10004D) was then added and samples were rotated for 1 hour at 4°C. The Dynabeads were then removed using a magnetic stand and the supernatants were transferred to new tubes. 5% of the total lysate was removed to be used as the input. 1 μg of rabbit anti-KDM1A antibody (Cell signaling, #2184S) was added and the samples were incubated overnight while rotating at 4°C. The following morning, 50 μl of Dynabeads were added and the samples were incubated for another 2 hours while rotating at 4°C. A magnetic stand was used to isolate the Dynabeads. The supernatant was removed and the Dynabeads were washed once in 1 mL of low salt immune complex wash buffer (20 mM Tris-HCl pH 8.0, 150 mM NaCl, 0.1% SDS, 1% TX-100, 2 mM EDTA) and incubated for 5 minutes while rotating at 4°C. The beads were again isolated using a magnetic stand. Following removal of the supernatant the Dynabeads were washed once in 1 mL of high salt immune complex wash buffer (20 mM Tris-HCl pH 8.0, 500 mM NaCl, 0.1% SDS, 1% TX-100, 2 mM EDTA) and incubated for 5 minutes while rotating at 4°C. The beads were again pelleted with a magnetic stand. Following removal of the supernatant, the Dynabeads were washed once in 1 mL of Lithium Chloride wash buffer (10 mM Tris-HCl pH 8.0, 250 mM LiCl, 1% SDS, 1% NP-40, 1 mM EDTA) and incubated for 5 minutes while rotating at 4°C. The beads were then washed twice in TE pH 8 at room temperature and the beads were then resuspended in 100 μl TE pH 8.

RNase A (Qiagen) was then added to a final concentration of 0.2 mg/mL and the samples were incubated for 30 minutes at 37°C. To reverse the crosslinking, proteinase K was then added to a final concentration of 0.8 mg/mL and the samples were incubated for 30 minutes at 42°C and then an additional 8 hours at 65°C. 300 μl QG buffer and 100 μl of isopropanol was then added to each sample and a magnetic stand was used to pellet the Dynabeads. The supernatant was then collected and DNA was isolated using QIAquick gel extraction kit (Qiagen) according to the manufacturer’s instructions.

The recovering DNA was analyzed by real-time qPCR. The DNA was quantified on a Bio-Rad T100 96-Well PCR Gradient Thermal Cycler with the SYBR Green kit (Bio-Rad, SsoAdvanced™ Universal SYBR® Green Supermix). The enrichment of DNA is shown as the percentage (%) input, which was calculated by determining the apparent IP efficiency at the ZFP36L1, REST, WWTR1 and YAP1 loci as the ratio of the amount of immunoprecipitated DNA normalized with input DNA. The following primers were used for qPCR:

ZFP36L1-1_Chip_For: ACACCTGGAGTCCCCTTGTT,

ZFP36L1-1_Chip_Rev: GATTTGGATCCAGGATTGCCT,

ZFP36L1-2_Chip_For: CCCCAAGGACTGCAAAATCC,

ZFP36L1-2_Chip_Rev: CCGGCCAGGTCTGATGAAA,

REST_Chip_For: GCAGGATCGGGTTACACAG,

REST_Chip_Rev: CTGACTCGGTGGAGCGTC,

WWTR1_Chip_For: GAAACTTGGTGGGGTGGCTA,

WWTR1_Chip_Rev: ACTGGTTCAGCAAGTCAGGG,

YAP1_Chip_For: GGAGTGTGCAGGAATGTAGCAA

CORL47 or NCI-H1876 cells were first infected with Cas9 and selected with blasticidin and then were superinfected with plentiguide-P2A-mCherry-puro^50^ encoding a puromycin resistance gene, and an sgRNA targeting the exon of ZFP36L1 (sgZFP36L1-1) or a non-targeting sgRNA (NOSITE-110). Puromycin-resistant cells, indicative of successful infection with the sgRNA lentiviruses, were counted using the Vi-Cell XR Cell and plated at 200,000 cells/mL for CORL47 cells or at 250,000 cells/ml for NCI-H1876 cells and then treated with ORY-1001 or CC-90011 and harvested after 7 days for cellular proliferation assays, RT-qPCR, immunoblot analysis, and RNA-sequencing experiments. For rescue experiments in Fig. S5, additional sgRNAs targeting exons of ZFP36L1 (sgZFP36L1-3,4,5, or 6) or a non-targeting sgRNA (C0111) were used.

### ZFP36L1 CRISPR Activation Experiments

CORL47 cells were first infected with lenti-dCas9-VP64_Blast (Addgene #61425) and selected with Blasticidin and then were superinfected with pXPR_502-Puro (Addgene #96923) lentivirus encoding a puromycin resistance gene, and an sgRNA targeting the promoter of *ZFP36L1* (sgZFP36L1-1 and sgZFP36L1-2) or a non-targeting sgRNA (C0111). Puromycin-resistant cells were selected with puromycin. Following generation of stable cell lines with endogenous CRISPR activation of ZFP36L1 confirmed by RT-qPCR, the cells were counted using the Vi-Cell XR and plated at 200,000 cells/mL for cell proliferation experiments, immunoblot analysis, and RNA-sequencing experiments.

### RNA-Sequencing

For the RNA sequencing experiment with NCI-H1876 Cas9 sgZFP36L1-1 or sgControl (NOSITE-110) cells, cells were counted and plated at 250,000 cells/ml and treated with ORY-1001 (1 nM or 100 nM) or DMSO and harvested after 7 days. For the RNA sequencing experiment in CORL47 cells with restored endogenous ZFP36L1 expression (see Method above), cells encoding sgZFP36L1-1, sgZFP36L1-2, or a non-targeting sgRNA were counted and plated at 200,000 cells/ml and harvested after 3 days. RNA was extracted using RNeasy mini kit including a DNase digestion step according to the manufacturer’s instructions and RNA sequencing was performed as described below.

Total RNA samples in each experiment were submitted to Novogene Inc. The libraries for RNA-seq are prepared using NEBNext Ultra II non-stranded kit. Paired end 150bp sequencing was performed on Novaseq6000 sequencer using S4 flow cell. Sequencing reads were mapped to the hg38 genome by STAR. Statistics for differentially expressed genes were calculated by DESeq2.

### RNA-Sequencing Analysis

For Gene Set Enrichment Analysis (GSEA), software was downloaded from the Gene Set Enrichment Analysis website [http://www.broad.mit.edu/gsea/downloads.jsp]. GSEA was performed using the “TOP 100 Neuroendocrine Gene Set”^47^ or “ASCL1 Target Genes DOWN”^51^ or Hallmarks Gene Sets. Gene Sets with an FDR q-value<0.25 were considered significant. For principal component analysis and cluster analysis, top 500 genes in terms of the largest standard deviation were subjected to principal component analysis using the prcomp function of R software. Clustering and heatmap were generated using the heatmap.2 function in the gplots package of R software.

### Analysis of Publicly Available RNA-Sequencing Data

To identify potential binding sites for LSD1 in target genes (as shown in Fig. S2), the Cistrome Data Browser was used. For the RNA expression in cancer cell lines in Fig. 3A, the RNA-sequencing data was downloaded from the Broad Institute Cancer Cell Line Encyclopedia.

For the RNA expression of ZFP36L1 in human lung tumors in Fig. 3B, the 81 SCLC RNA-Seq data was previously published ^4^ and publicly available on the cBioPortal^49^. The 747 lung adenocarcinoma samples were from TCGA and publicly available on the cBioPortal^49^. The ZFP36L1 RNA expression for each sample was normalized to the ACTB RNA for that sample. The MHC class I high vs. low human SCLC RNA-sequencing data set was previously described^37^.

For the analysis of RNA-Seq data from SCLC PDX models, a previously published RNA-sequencing data set, where SCLC PDX models were treated *in vivo* and *ex vivo* with ORY-1001 or the vehicle control, was used^7^. The RNA-Seq fastq files were downloaded from the GEO (accession number GSE103097)^7^. Sequenced reads were aligned to both hg19 and mm10 genomes using STAR. Then the alignment files were processed with bamcmp package^52^ to distinguish between human and mouse reads (“human only” and “human better” reads were retained for further analysis). The featureCounts algorithm^53^ was used to quantify transcript abundance. To calculate the enrichment of hits from the CRISPR/Cas9 ORY-1001/KDM5-C70 screen, AUCell package^54^ was utilized. This package has been originally developed for measuring enrichment of given signatures in scRNA-Seq data and it was adapted for our purposes with minor modifications. For all drug-treated CRISPR/Cas9 screens, hits were included if they had a q-value less than 0.25 from the STARS analysis. For the DMSO-treated CRISPR/Cas9 screen, hits were included that had a p-value less than 0.05 as there were very few hits with a q-value of less than 0.25.

To generate heatmaps of relative changes in the SCLC PDX models after ORY-1001 treatment^7^, the list of genes for each arm of the CRISPR/Cas9 screen was further curated based on the statistical cutoffs listed above. First, genes were ranked in each model and the relative change in ranking was calculated based on the following formula: Relative Change=(Rank Before ORY-1001 Treatment-Rank After ORY-1001 Treatment)/Rank Before ORY-1001 Treatment. Then, the heatmap function in R was used to generate heatmaps and cluster genes/samples based on the relative change.

### Tumor Micro Array (TMA) construction

Tumor cores were obtained from embedded PDX tumors tissue in donor blocks using a 1 mm biopsy punch needle (IHC World; W125-0) then embedded/inserted into paraffin recipient block/negative mold (IHC World; 10*17 Quick Ray mold IW-UM01-1). Empty slots were filled with blank paraffin cores. All cores were gently tamped down using biopsy punch needle then the entire block was placed face down on glass slides and heated to 50° C for 2 hours to merge donor cores with the recipient block. Blocks and glass slides were then placed on ice for 30 minutes before sectioning.

### Immunohistochemistry (IHC)

Mouse tumor samples used for IHC experiments were from our previously described CRISPR-based SCLC genetically-engineered mouse model^15^. These samples were generated by injecting adenoviruses encoding Cre-recombinase and sgRNAs targeting *Rb1, Trp53, Rbl2,* and a non-targeting sgRNA (sgControl RPP) into the lungs of homozygous Lox-stop-lox (LSL) Cas9 B6J mice (Jackson No. 026175). Immunohistochemical staining for Zfp36l1 and Ascl1 was performed by hand. For immunohistochemistry experiments, 4 µm thick tissue sections were prepared and left to air-dry overnight. Slides were baked in an Isotemp Oven (Fisher Scientific) for 30 minutes at 60°C to melt excess paraffin. For antigen retrieval, slides were heated with a DC2002 Decloaking Chamber (Biocare Medical, Pacheco, CA) to 125°C for 30 seconds and 90°C for 10 seconds in citrate (pH = 6.0, Thermo Scientific cat. no. Ap-9003-500) for Ascl1 and Zfp36l1 staining. All sections were incubated with peroxidase (Dako cat. no. S2003, Carpinteria, CA) and protein (Dako cat. no. X0909) blocking reagents for 5 minutes each. Sections were then incubated with rabbit anti-TISB (ZFP36L1) antibody (Sigma #SAB4502867; 1:100) or rabbit monoclonal anti-Ascl1 antibody (Abcam #ab211327; 1:100) diluted in Dako Diluent with Background Reducing Components (Dako #S3022) for 60 minutes at room temperature. Following primary incubation, sections were incubated with Envision+System-HRP Labelled Polymer Anti-Rabbit (Dako #K4003) for 30 minutes. All sections were developed using the DAB chromogen kit (Dako #K3468) and counterstained with hematoxylin, dehydrated in graded ethanol and xylene, mounted, and coverslipped. To establish specificity of the IHC assay for Zfp36l1, we compared PC9 cells infected with an sgRNA targeting ZFP36L1 or a non-targeting sgRNA control using PC9 isogenic cells shown in Fig. 4C. Our Ascl1 IHC assay was performed and validated as previously described^15^. Whole slides images were acquired using an Aperio AT scanner (Leica). For each stain, an intensity threshold was defined using the Multiplex IHC v2.3.4 algorithm with Halo (Indica Labs). Pseudo-immunofluorescence images were obtained by deconvoluting the immunohistochemistry images using the Deconvolution v1.0.3 algorithm with Halo for better visualization.

### RNA-Binding Protein Immunoprecipitation (RIP)

For experiments in CORL47 cells with exogenous constitutive ZFP36L1 expression, PLX304-CMV-ZFP36L1WT-V5-BLAST or PLX304-CMV-EV-V5-BLAST lentiviruses were infected into CORL47 cells and selected with blasticidin. After selection and confirmation of expression of ZFP36L1 via immunoblot analysis, the cells were counted on day 0 using a Vi-Cell XR Cell Counter and plated at 2 × 10^6^ cells in 10 mLs of complete media. For experiments in CORL47 cells with exogenous DOX-On inducible ZFP36L1 expression, DOX-On inducible pTripz-ZFP36L1WT-FLAG-NEO, pTripz-ZFP36L1MUT-FLAG-NEO, or pTripz-EV-FLAG-NEO lentiviruses were infected into CORL47 cells and selected with G418. After selection and confirmation of expression of ZFP36L1 via immunoblot analysis in the presence of DOX, all cell lines were counted on day 0 using a Vi-Cell XR Cell Counter and plated at 2 × 10^6^ cells in 10 mLs of complete media for 3 days. For the DOX-On pTripZ experiments, cells were grown in the presence or absence or Doxycycline (DOX) (1 μg/mL) as indicated.

3 days after plating, the cells were collected, centrifuged, and the supernatant was removed. The cells were washed once in cold PBS and again centrifuged for 5 minutes at 1500 rpm at 4°C. The cell pellets were then resuspended in 500 μL of cold PBS, and then 50 μL was removed for input RNA (10% of total cell lysate) and 25 μL for input protein (5% of total cell lysate), and the other 425 μL was used for IP. The cells were centrifuged again and the supernatant was removed. The cell pellets were resuspended in 200 μL of RIP lysis buffer (20 mM Tris-HCl pH 7.5, 100 mM NaCl, 0.5 % NP-40, 5 mM MgCl2, protease inhibitor cocktail (Complete, Roche Applied Science, 11836153001), phosphatase inhibitors (PhosSTOP Sigma #04906837001) and RNase inhibitor (RNaseOUT, Thermofisher #10777019). The lysate was mixed up and down by pipetting until the mixture was homogeneous. The lysate was incubated on ice for 10 minutes and then centrifuged for 10 minutes at 13,500 rpm at 4°C. The lysate was then aliquoted at equal volumes (100 μL and 100 μL) to each dynabeads-antibody complex containing the antibody of interest or the isotype-matched IgG negative control.

To conjugate the magnetic beads with antibody, Dynabeads Protein G (ThermoFisher #10004D) were first resuspended by pipetting and then 25 μL/reaction of Dynabead suspension was transferred to each eppendorf tube. Then 0.5 mLs of RIP wash buffer (50 mM Tris-HCl pH7.5, 150 mM NaCl, 1 mM MgCl2, 0.05 % NP-40) was added to each tube containing Dynabeads and vortexed briefly. The tubes were then placed on the magnet (DynaMag™-2 Magnet, ThermoFisher #12321D) to isolate the Dynabeads and the supernatant was discarded. The eppendorf tube was then removed from the magnet and RIP wash buffer was added to wash. This was repeated 3 times. Then the Dynabeads were resuspended in 100 μL of RIP wash buffer and 2.5 μL (2.5 μg) of mouse anti-FLAG (Sigma, clone M2, #F1804-1MG) antibody or isotype-matched control IgG1 (Cell Signaling #5415S) to the tube for the FLAG IPs, or 5 μL of rabbit anti-V5 (Cell Signaling #13202) or the isotype-matched IgG control (Cell Signaling #3900) for the V5 IP. The antibody-bead mixtures were then incubated while rotating for 30 minutes at room temperature to conjugate the antibodies with the Dynabeads. Then 100 μL of protein lysate (∼1 mg of protein determined in optimization pilot experiments performed to determine cell plating density) was mixed with conjugated antibody-Dynabead complex (100 μL) and 900 μL of RIP immunoprecipitation buffer (860 μL RIP wash buffer, 35 μL of 0.5 M EDTA, 5 μL RNase inhibitor) for total volume of 1.1 mL. The reaction was incubated overnight with rotating at 4 °C.

The following morning, the immunoprecipitation Eppendorf tubes were briefly centrifuged and then placed on the magnet and the supernatant was discarded. The tubes were removed from the magnet, and then 0.4 mL of cold RIP wash buffer was added. This was repeated 3 times. During the final wash, 40 μL of a 400 μL total volume was removed to test the IP efficiency by placing the 40 μL on the magnet to remove the RIP wash buffer and the Dynabeads were resuspended in 30 μL 2 x SDS-PAGE loading buffer and the samples were boiled at 95 °C for 10 minutes. The samples were then run on a 10% SDS-PAGE with the input protein samples described above.

For the remaining 360 μL of the IP, the Dynabeads were isolated by the magnet and then resuspended in 150 μL proteinase K buffer [130 μL RIP wash buffer, 15 μL 10 % SDS, 5 μL 40 mg/mL proteinase K (Invitrogen #25530-015) and incubated at 55 °C for 30 minutes with shaking. The tube was then spun down briefly and placed on the magnet. The supernatant was then transferred into a new Eppendorf tube for RNA isolation.

For RNA extraction the Quick-RNA Miniprep Kit (ZYMO Research, # R1054) was used according to the manufacturer’s instructions with the following modifications. 450 μL cell lysis buffer from the RNA kit was added to the 150 μL of the IP (or 550 μL cell lysis buffer was added to 50 μL of the RNA input described above) so that the final volume was 600 μL. At the elution step, 20 μL RNase/DNase free water was added and 8 μL of eluted RNA was used for cDNA library synthesis.

A cDNA library was synthesized using iScript Reverse Transcription Supermix for RT-qPCR (Biorad # 1708841) according to the manufacturer’s instructions. qPCR was performed using the SYBR Green PCR Kit (Bio-Rad, SsoAdvanced™ Universal SYBR® Green Supermix) according to the manufacturer’s instructions. Signals were detected using the Bio-Rad T100 96- Well PCR Gradient Thermal Cycler. The ÄÄC_T_ Method was used to analyze data. The expression level of each gene was normalized to the percentage (%) input. The primers for qPCR are:

SOX2-For_ TACAGCATGATGCAGGACCA,

SOX2-Rev_CCGTTCATGTAGGTCTGCGA,

INSM1-For_TTACGCGGCACATCAACAAG,

INSM1-Rev_TCTTTGTGGGTCTCCAAGCG,

CDK2-For_CATCTTTGCTGAGATGGTGACT,

CDK2-Rev_ACTTGGCTTGTAATCAGGCAT,

E2F1-For_CCGGGGAATGAAGGTGAACA,

E2F1-Rev_GAGCAAAAGGGCCGAAAGTG,

GAPDH-For_ TCGGAGTCAACGGATTTGGTC,

### 3’ UTR Luciferase Stability Assay

For the 3’UTR luciferase assay, first 293T cells with exogenous constitutive ZFP36L1 expression were established. The PLX304-CMV-ZFP36L1WT-V5-BLAST, or PLX304-CMV- ZFP36L1MUT-V5-BLAST, or PLX304-CMV-EV-V5-BLAST lentiviruses were infected into 293T cells and selected with blasticidin and confirmed to express ZFP36L1 via immunoblot analysis. Then all cell lines were counted on day 0 using a Vi-Cell XR Cell Counter and plated at 1 × 10^5^ cells in 0.5 mL in a 24-well tissue culture plate in DMEM media with 10% FBS without Penicillin and Streptomycin and incubated overnight. The following morning, the cells were transfected with pLightSwitch-3’UTR control vector, the pLightSwitch-hSOX2-3’UTR, or the pLightSwitch- hINSM1-3’UTR plasmid using Lipofectamine 2000 reagent (Invitrogen™, Cat #: 11668019) according to the manufacturer’s instructions. 16 to 18 hours later, luciferase activity was measured using LightSwitch™ Luciferase Assay Kit (SwitchGear Genomics, LS010) and Infinite 200 PRO microplate plate reader. Relative luciferase was normalized with cell number, and then normalized with pLX304-EV group.

### siRNA Experiments

Nucleofection was performed using the SF 4D-Nucleofector X kits (Lonza # V4XC-2032). For transfection, NCI-H1876 and CORL47 (1 × 10^6^ cells) were resuspended in 20 μL of SF 4D-Nucleofector X solution and transferred into a the 16-well Nucleocuvette strip. 2 µM of siRNA was added and electroporation was performed using the CM-137 program (C1). After transfection, 70 µL of pre-warmed culture medium was gently added to each sample and incubated for 10 minutes at 37 C. Transfected cells in suspension were then gently transferred dropwise into 6-well tissue culture-treated plates containing 2 mL pre-warmed complete media. Cells were then cultured for 72 hours and 96 hours and harvested for immunoblot analysis. The following siRNAs were used: on-target plus human SOX2 siRNA SMARTpool (Horizon, Catalog ID:L-011778-00), on-target plus human INSM1 siRNA SMARTpool (Horizon, Catalog ID:L- 006535-00), and on-target plus non-targeting siRNA (Horizon, Catalog ID:

### Statistical Analysis

For the CRISPR/Cas9 screen in Figure 1, the data was analyzed using the STARS and Hypergeometric analysis algorithms on the GPP portal (https://portals.broadinstitute.org/gpp/screener/) and genes were considered “hits” if their q-value was <0.25 using the STARS analysis. For the RNA sequencing experiments and GSEA analysis in Figure 4, statistical significance with calculated using FDR corrected for multiple hypothesis testing where q-value of <0.25 is considered statistically significant.

For all other experiments, statistical significance was calculated using unpaired, two- tailed Students t-test. *p*-values were considered statistically significant if the *p*-value was <0.05. For all figures, * indicates *p*-value <0.05, ** indicates *p*-value <0.01, *** indicates and *p*-value <0.001, **** indicates and *p*-value <0.0001. Error bars represent SEM unless otherwise indicated.

