## Supplemental Figures and Legends for "Regulation of Neuroendocrine Plasticity by the RNA-Binding Protein ZFP36L1"

**Supplementary Data**

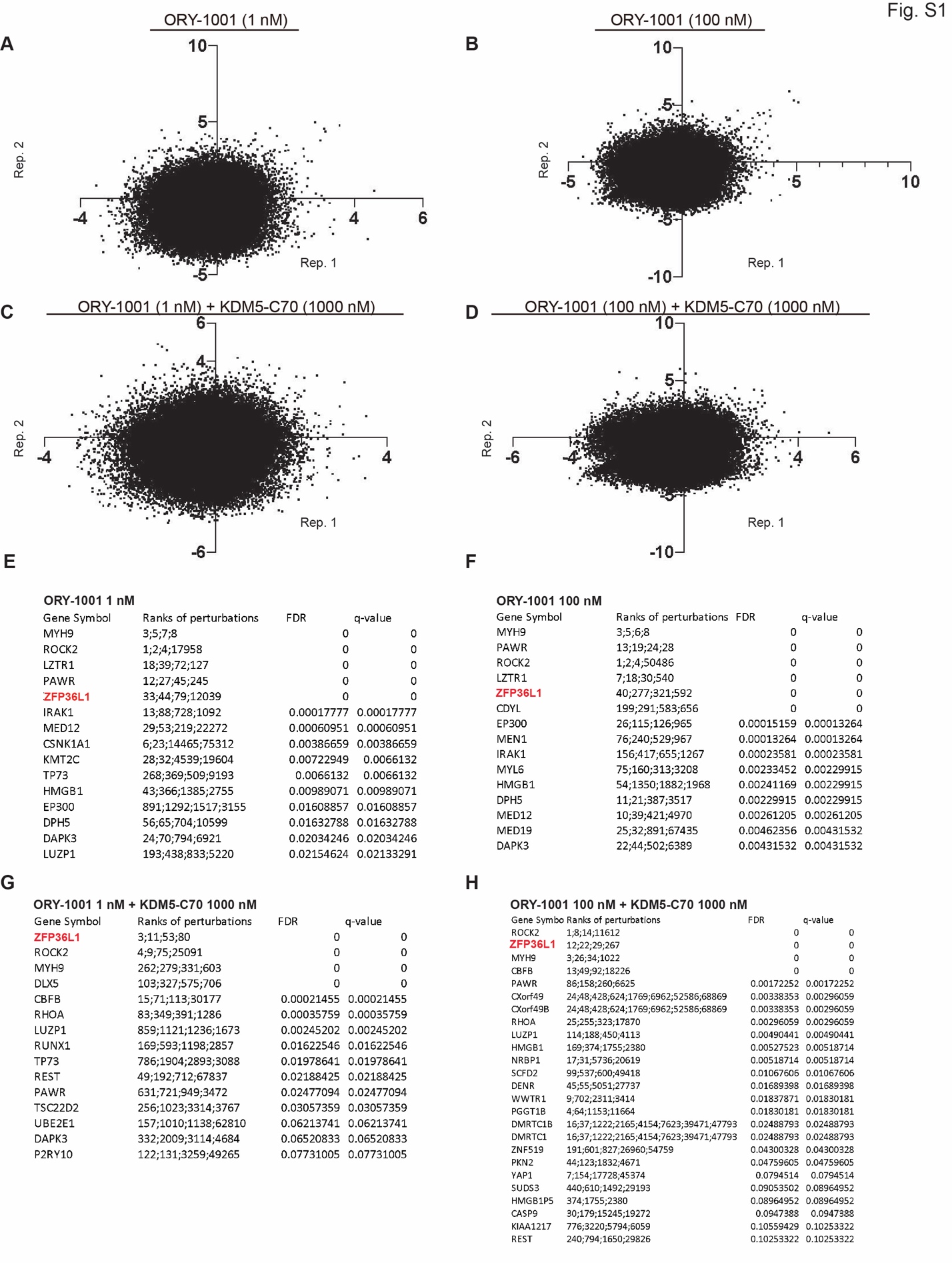

**Extended Data Fig. S1. CRISPR/Cas9 Positive Selection Screen Identifies Genes Required for LSD1 Inhibitor Sensitivity in Small Cell Lung Cancer.** (**A-D**) Log-fold change (LFC) of sgRNAs on day 35 relative to the early timepoint (day 11) of screen replicate 1 compared to screen replicate 2 of the screens treated with ORY-1001 (1 nM) (**A**), ORY-1001 (100 nM) (**B**), ORY-1001 (1 nM) + KDM5-C70 (1000 nM) (**C**), or ORY-1001 (100 nM) + KDM5-C70 (1000 nM) (**D**). n=2 biological replicates. (**E-H**) STARS analysis from the positive-selection CRISPR/Cas9 screen on day 35 relative to the early timepoint prior to drug treatment (day 11) of NCI-H1876 Cas9 cells infected with the Brunello sgRNA library and then treated with ORY-1001 (1 nM) (**E**), ORY-1001 (100 nM) (**F**), ORY-1001 (1 nM) + KDM5-C70 (1000 nM) (**G**), or ORY-1001 (100 nM) + KDM5-C70 (1000 nM) (**H**). n=2 biological replicates.

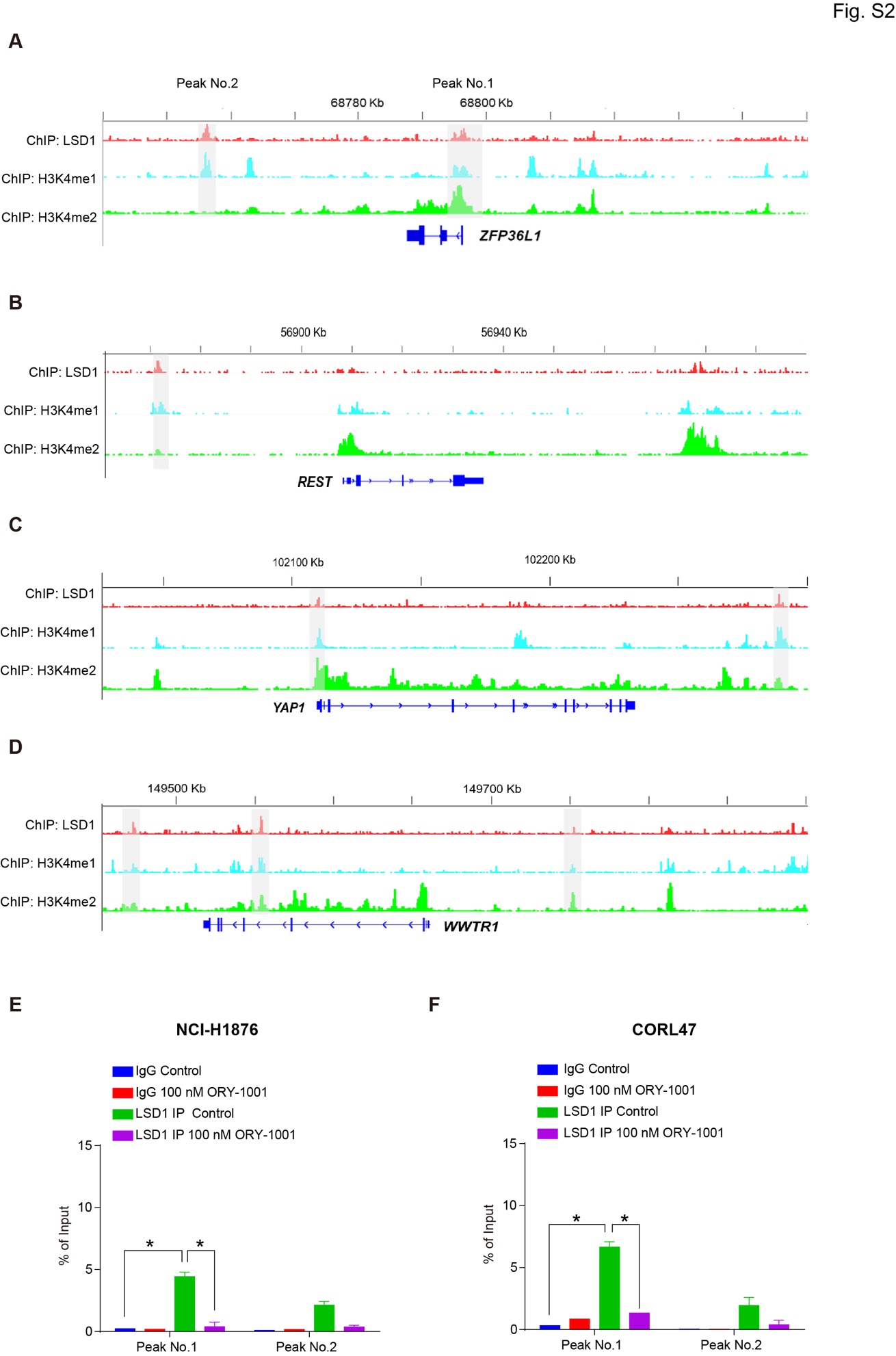

**Extended Data Fig. S2. LSD1 Binds ZFP36L1, REST, YAP1 and WWTR1.**

(**A-D**) Read density tracks, visualized using cistrome.db database, of normalized ChIP-seq for LSD1 (red), H3K4me1 (blue) and H3K4me2 (green) in SH-SY5Y neuroblastoma cells. Shaded area highlights region of ZFP36L1 (**A**), REST (**B**), YAP1 (**C**) and WWTR1 (**D**) with LSD1/H3K4me1/H3K4me2 binding. (**E and F**) ChIP-qPCR of NCI-H1876 (**E**) and CORL47 (**F**) cells first treated with ORY-1001 100 nM or DMSO for 6 days before performing immunoprecipitation (IP) for LSD1 followed by qPCR of ZFP36L1 of Peak No. 1 and Peak No. 2 as indicated in S2A. Peak No. 1 is the peak used in Fig. 2H&I. n=2 biological replicates. For all panels, *=p<0.05.

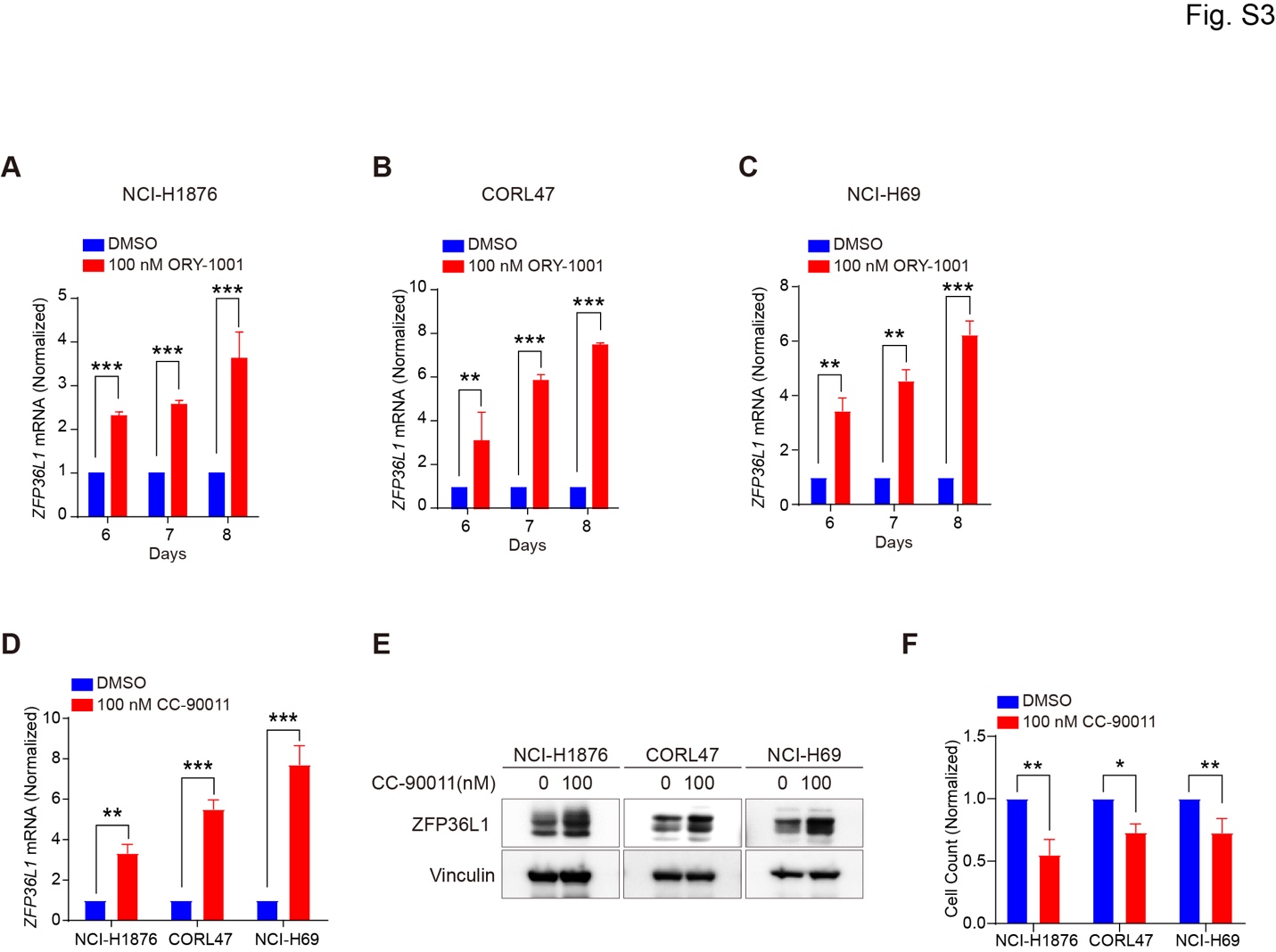

**Extended Data Fig. S3. ZFP36L1 is Induced by LSD1 Inhibitors.**

(**A-C**) RT-qPCR of NCI-H1876 (**A**), CORL47 (**B**), and NCI-H69 cells (**C**) treated with ORY-1001 (100 nM) for 6, 7 and 8 days. n=2 biological replicates. (**D-F**) RT-qPCR (**D**), immunoblot analysis (**E**), and cell counts (**F**) of NCI-H1876, CORL47, and NCI-H69 cells treated with the LSD1 inhibitor CC-90011 (100 nM) for 7 days. For D, n=4 biological replicates. For F, n=2 biological replicates. For all panels, *=p<0.05, **=p<0.01, ***=p<0.001.

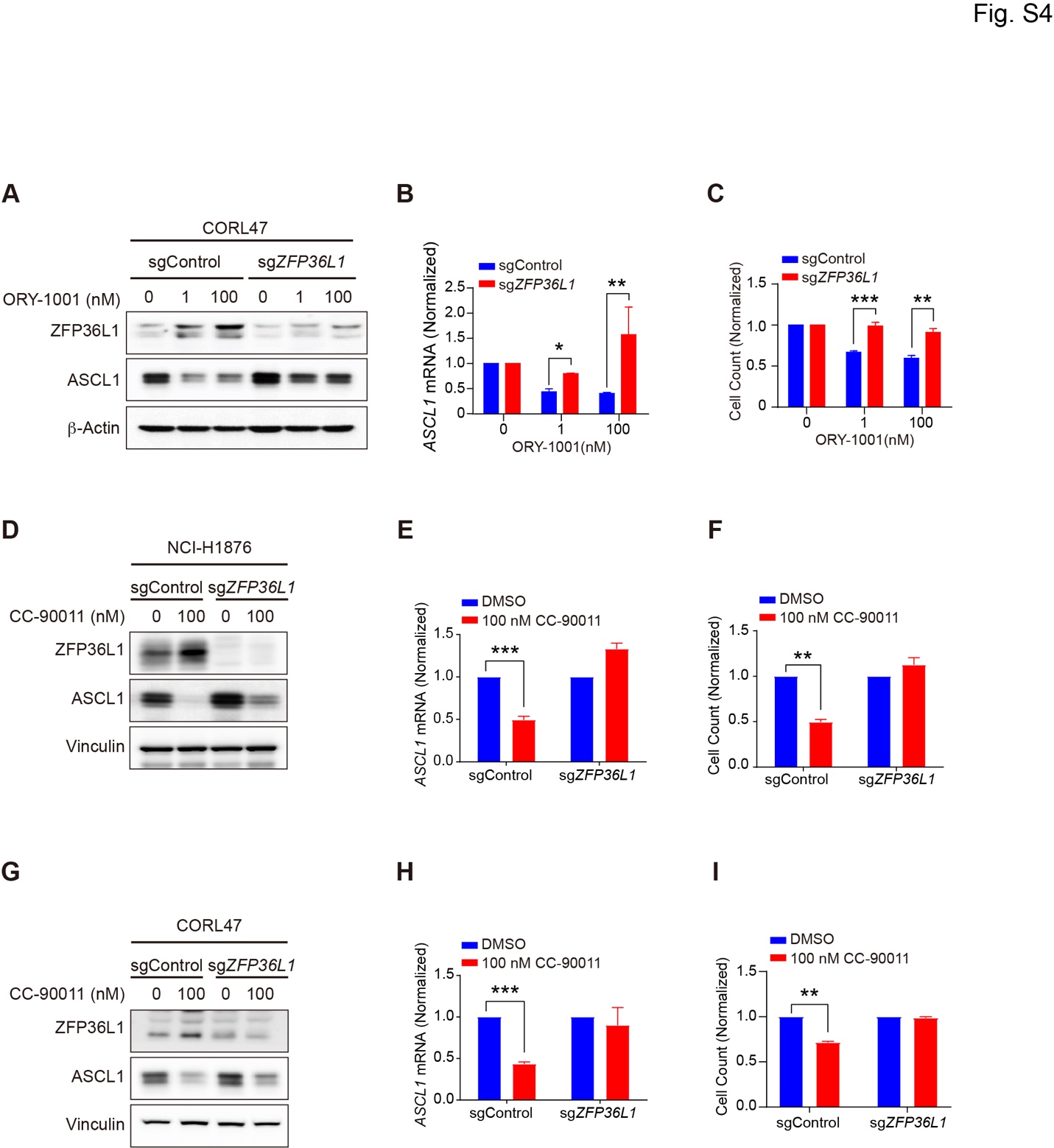

**Extended Data Fig. S4. LSD1 Inhibitors Block Neuroendocrine Differentiation and Proliferation in Small Cell Lung Cancer Through a ZFP36L1-Dependent Mechanism**

(**A-C**) Immunoblot analysis (**A**), RT-qPCR (**B**), and cell counts (**C**) of CORL47 Cas9 cells infected with lentiviruses encoding an sgRNA targeting ZFP36L1 (sgZFP36L1) or a non-targeting sgRNA (sgControl) and then treated with ORY-1001 (1 nM and 100 nM) or DMSO for 7 days. For B, n=2 biological replicates. For C, n=4 biological replicates. (**D-I**) Immunoblot analysis (**D and G**), RT-qPCR (**E and H**) and quantitation of cell counts (**F and I**) of NCI-H1876 Cas9 cells (**D-F**) and CORL47 Cas9 cells (**G-I**) first infected with sgZFP36L1 or sgControl lentiviruses and then treated with CC-90011 (100 nM) or DMSO for 7 days. For E, n=3 biological replicates. For F, n=2 biological replicates. For H, n=4 biological replicates. For I, n=2 biological replicates. For all panels, *=p<0.05, **=p<0.01, ***=p<0.001.

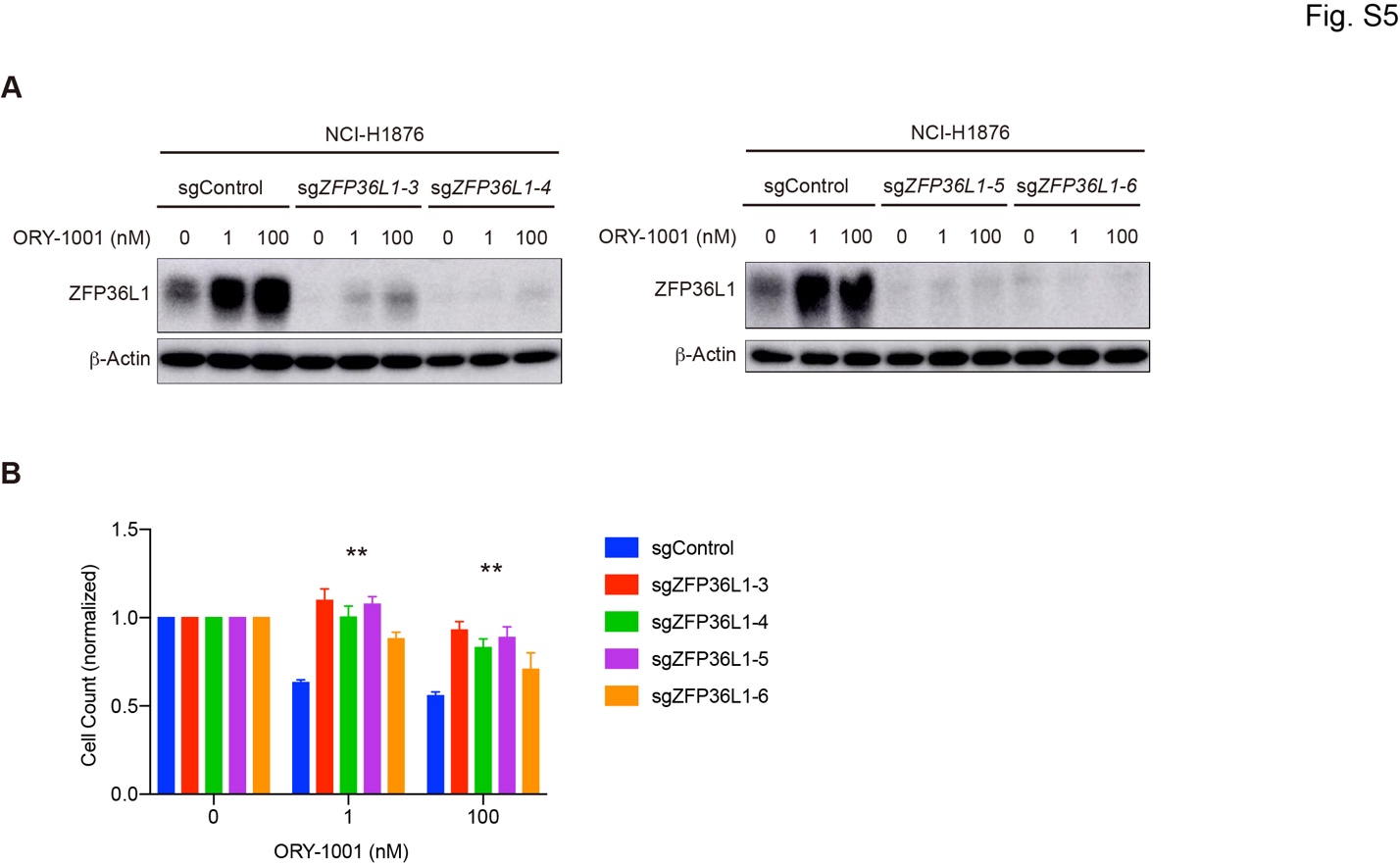

**Extended Data Fig. S5. ZFP36L1 is Required for the Anti-Proliferative Effects of ORY-1001 in NCI-H1876 Cells**

Immunoblot analysis (**A**) and quantitation of cell counts (**B**) of NCI-H1876 Cas9 cells infected lentiviruses encoding 4 independent sgRNAs targeting ZFP36L1 (labeled as 3-6) or a non-targeting sgRNA (sgControl) and then treated with ORY-1001 (1 nM and 100 nM) or DMSO for 7 days. n=4 biological replicates. **=p<0.01 for all ZFP36L1 sgRNAs compared to sgControl at both ORY-1001 concentrations except for sgZFP36L1-6 vs. sgControl at 100 nM where p=0.16.

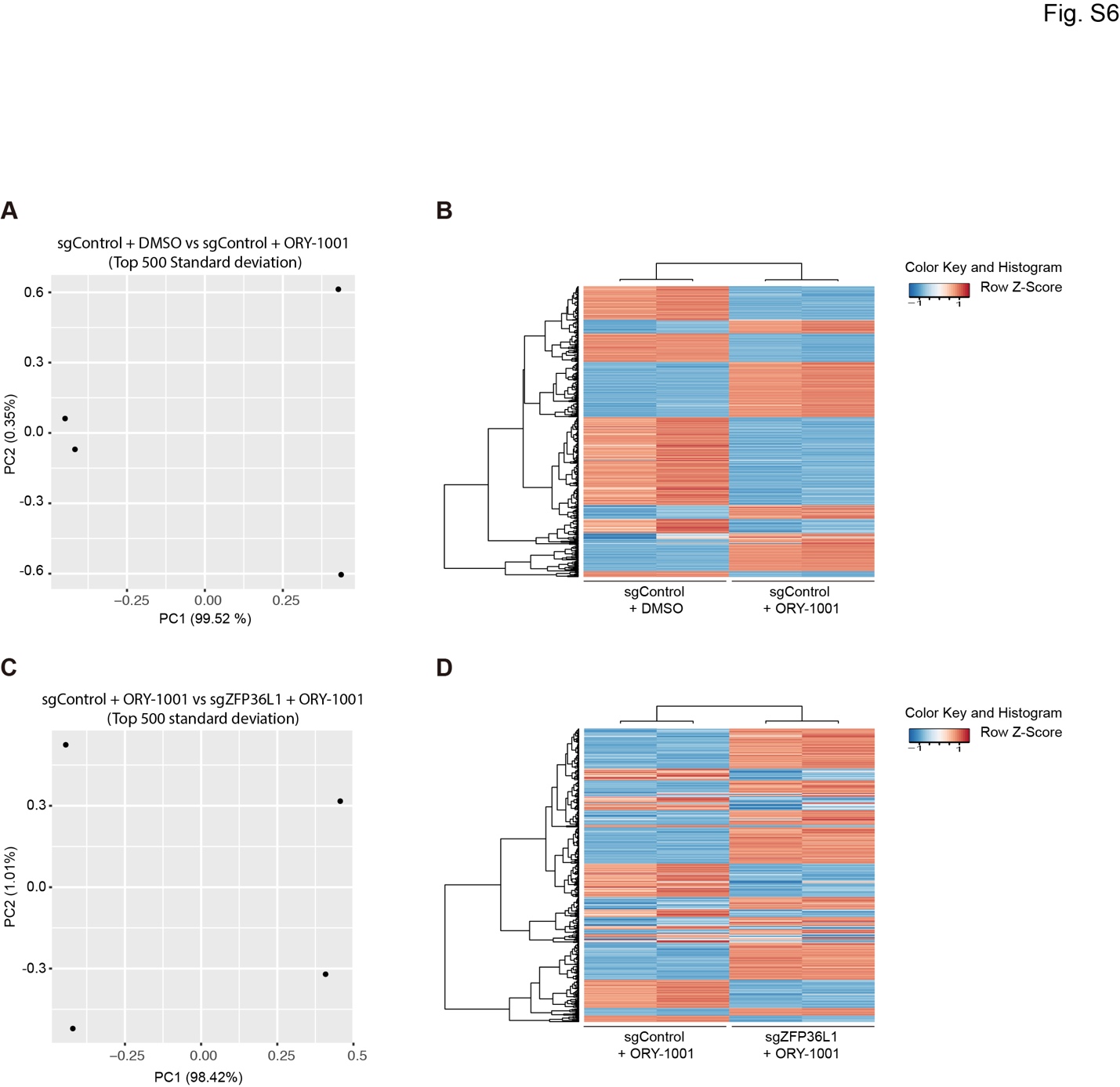

**Extended Data Fig. S6. RNA-Sequencing Analysis of sgZFP36L1 vs. sgControl NCI-H1876 Cells Treated with ORY-1001**

(**A**) Principal component analysis (PCA) of gene expression from RNA-seq data from Fig. 4F of NCI-H1876 sgControl cells treated with ORY-1001 (1 nM) or DMSO. (**B**) Unsupervised hierarchical clustering heat map of top 500 upregulated and top 500 downregulated genes in NCI-H1876 sgControl DMSO vs. sgControl ORY-1001 (1 nM) cells from the RNA-Seq experiment in A and Fig. 4F. (**C**) PCA of gene expression from RNA-seq data from Fig. 4H of NCI-H1876 sgControl cells treated with ORY-1001 (1 nM) or sgZFP36L1 cells treated with ORY-1001 (1 nM). (**D**)Unsupervised hierarchical clustering heat map of top 500 upregulated and top 500 downregulated genes in NCI-H1876 sgControl ORY-1001 (1 nM) vs. sgZFP36L1 ORY-1001 (1 nM) cells from the RNA-Seq experiment in C and Fig. 4H. For B and D, the red to blue color scale indicates FPKM values from large to small. n=2 biological replicates.

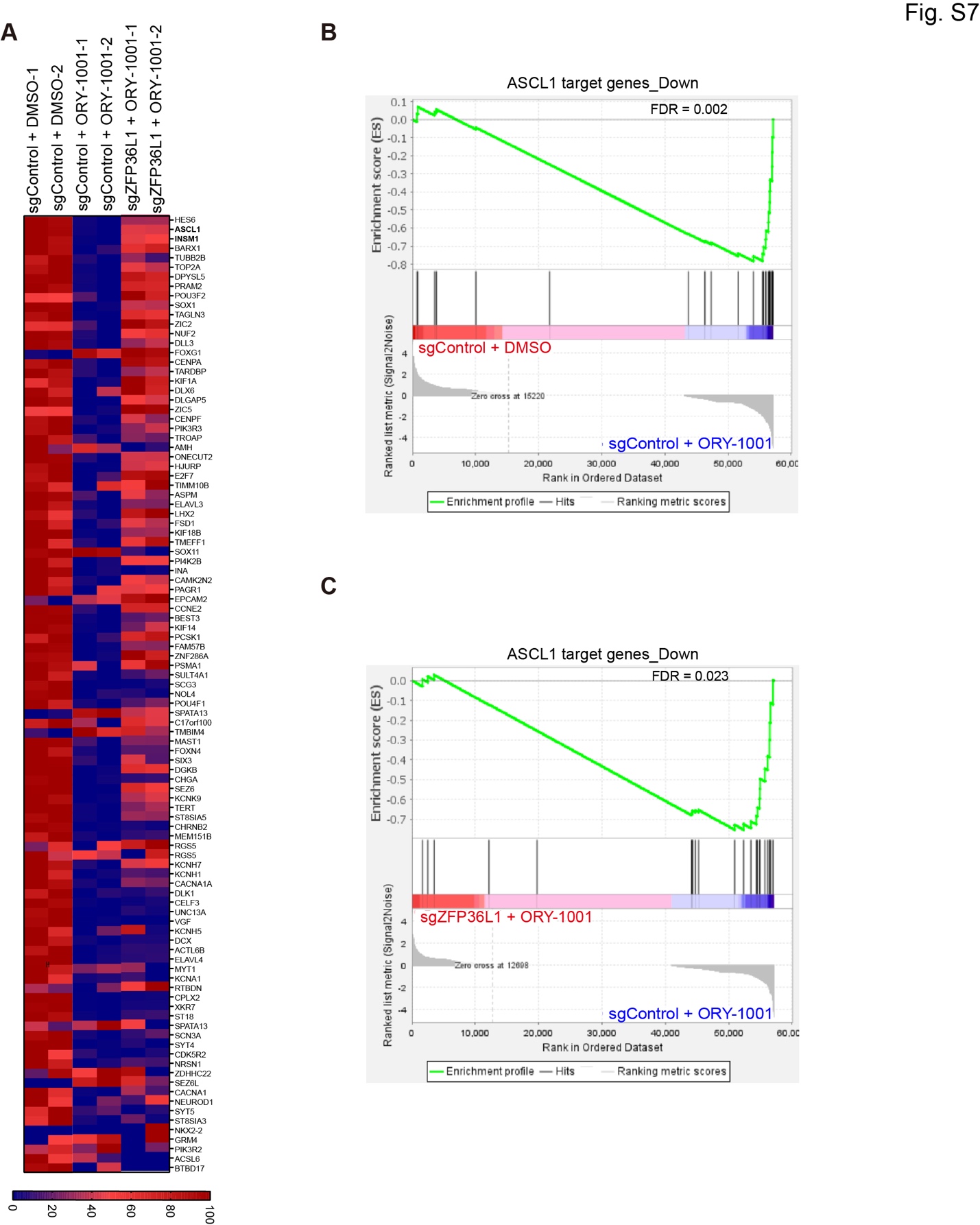

**Extended Data Fig. S7. Gene Set Enrichment Analysis of RNA-Seq Data of sgZFP36L1 vs. sgControl NCI-H1876 Cells Treated with ORY-1001**

(**A**) Heatmap of the changes in the Top 100 Neuroendocrine Genes from the RNA-seq experiment in Fig. 4F-I of NCI-H1876 cells with the perturbations indicated. Red denotes genes with high expression, and blue denotes genes with low expression. (**B and C**) Gene set enrichment analysis (GSEA) of RNA-seq data in Fig. 4F (**B**) or 4H (**C**) of the ASCL1 Target Genes Down gene set. FDR q-values are indicated. n=2 biological replicates.

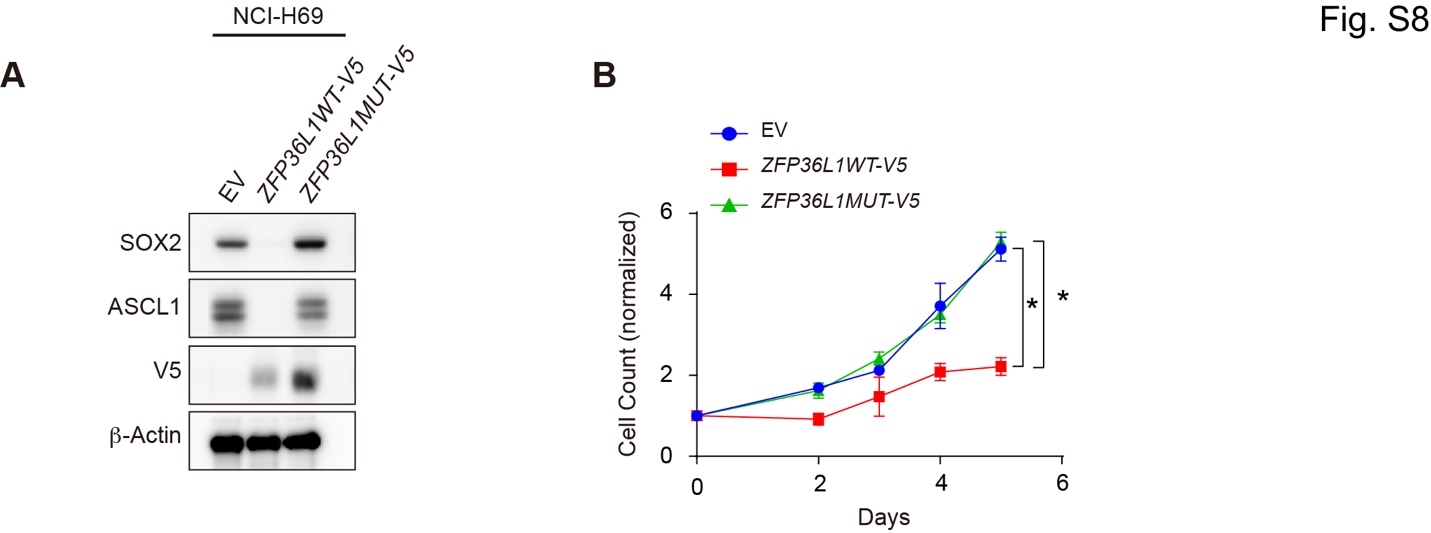
 **Extended Data Fig. S8. The mRNA-binding activity of ZFP36L1 is required to block neuroendocrine differentiation and cellular proliferation in NCI-H69 Cells.**

(**A**) Immunoblot analysis of NCI-H69 cells stably infected with ZFP36L1WT-V5, the ZFP36L1 mRNA-binding mutant-V5 (ZFP36L1MUT-V5), or the corresponding empty vector (EV). (**B**) Proliferation assays of the cells in A. n=3 biological replicates. *=p<0.05.

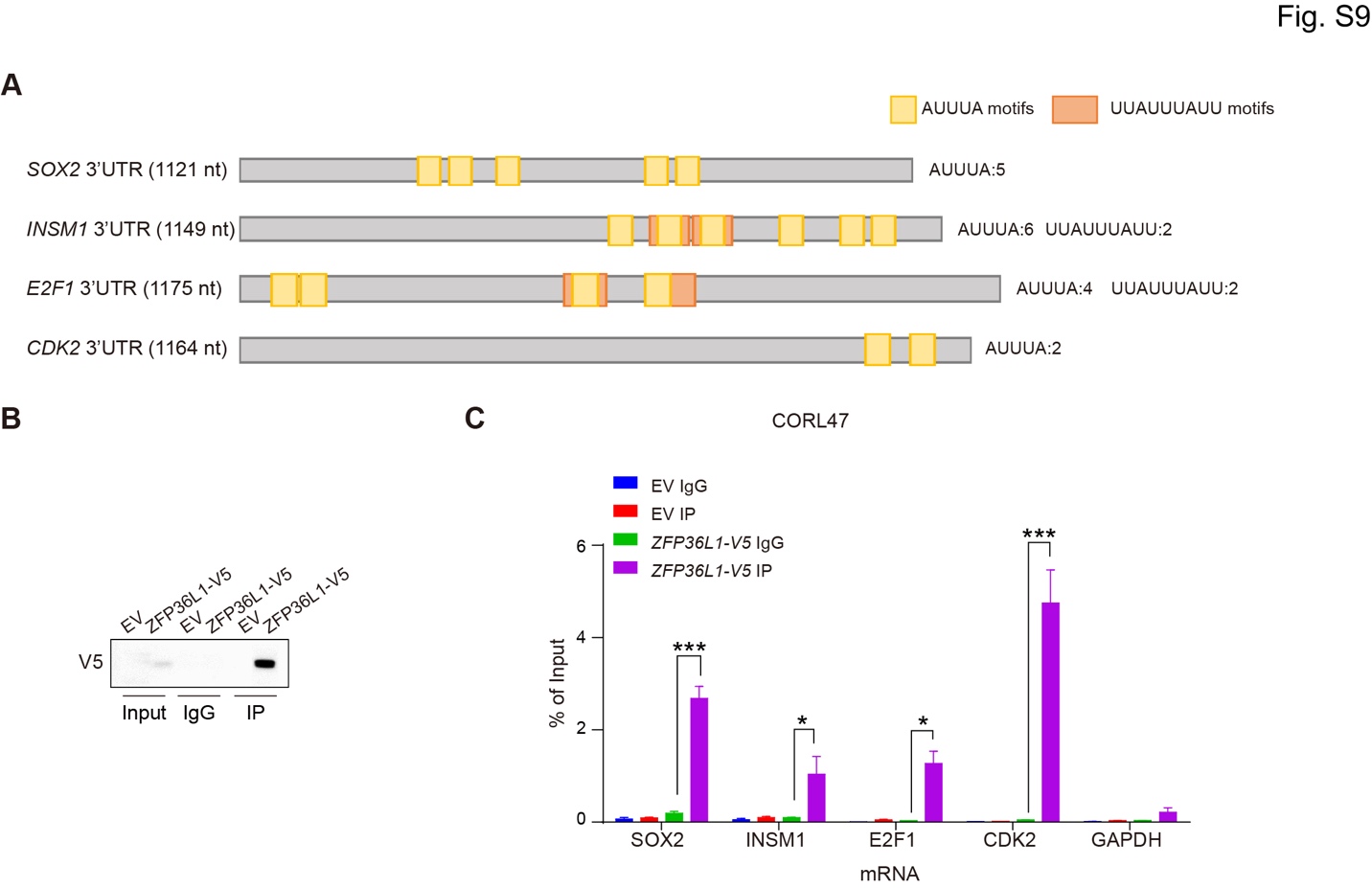

**Extended Data Fig. S9. ZFP36L1 Binds SOX2, INSM1, E2F1, and CDK2 mRNAs.**

(**A**) Schematic of the ZFP36L1 canonical binding sites (AU-rich elements=AREs) in the 3’UTR’s of SOX2, INSM1, E2F1, and CDK2. The number of ARE’s are indicated. Yellow is AUUUA motifs and orange is UUAUUUAUU motifs. (**B**) Immunoblot analysis after immunoprecipitation (IP) of ZFP36L1WT-V5, ZFP36L1MUT-V5, or EV from CORL47 cells relative to input. (**C**) mRNA quantitation relative input after RT-qPCR from IP experiments in B with primers specific to SOX2, INSM1, E2F1, CDK2, and GAPDH. n=2 biological replicates. *=p<0.05, ***=p<0.001.

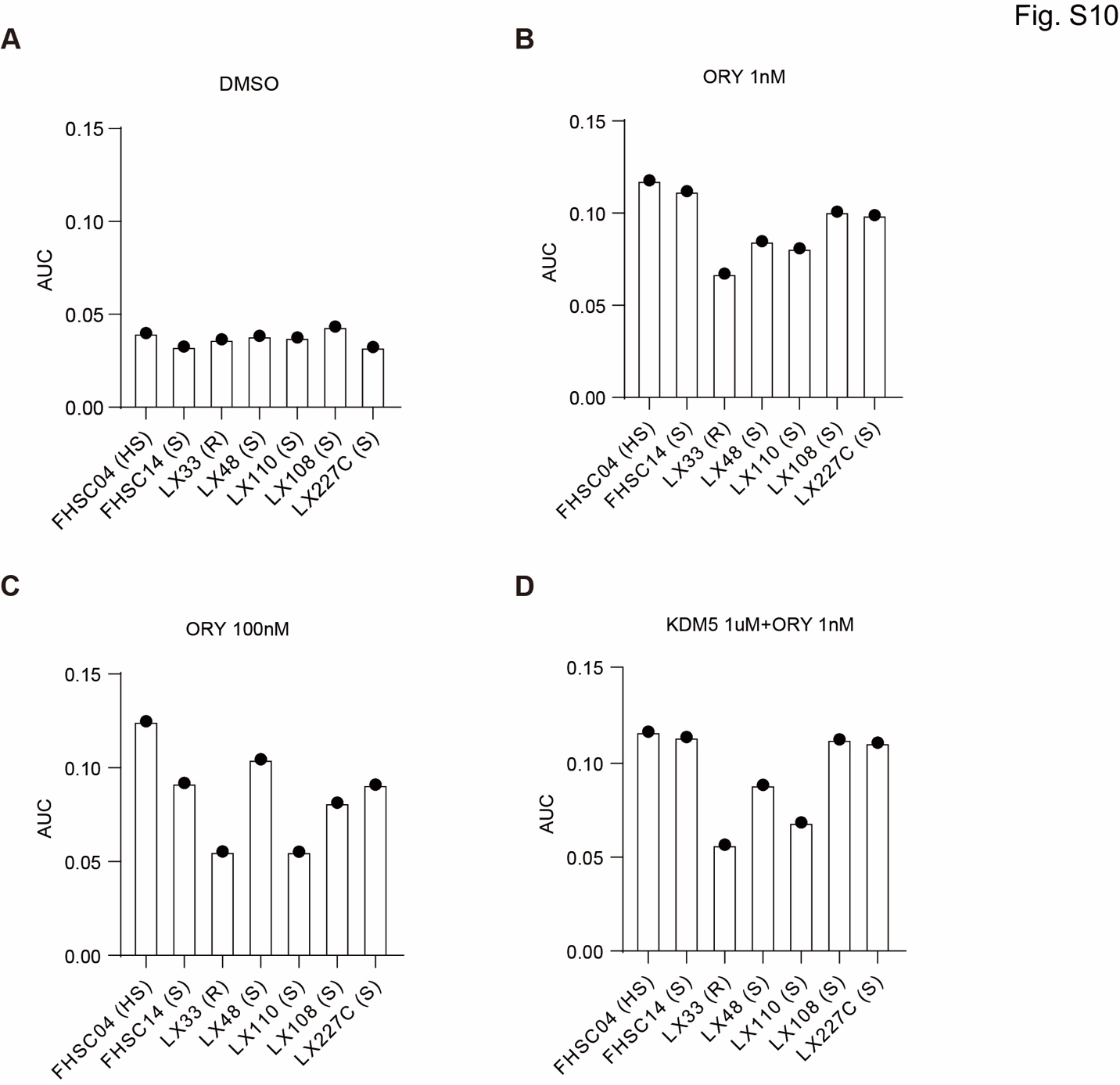

**Extended Data Fig. S10. Enrichment Analysis of Correlation of Expression of Hits from our CRISPR/Cas9 ORY-1001/KDM5-C70 Resistance Screen with ORY-1001 Sensitivity.**

(**A-D**)Area under the curve (AUC) enrichment analysis of hits from our ORY-1001/KDM5-C70 resistance screen (see Fig. 1C-F) of RNA-seq data from untreated patient-derived xenograft (PDX) models of SCLC. Hits were considered significant and were included in the AUC enrichment analysis if their q-value was less than 0.25 for ORY-1001 1 nM (**B**), ORY-1001 100 nM (**C**), and ORY-1001 (1 nM) + KDM5-C70 (1000 nM) (**D**). Hits were considered significant in the DMSO arm with p-values less than 0.05. A less strigent cut-off was used in the DMSO arm because there were very few hits with q-value less than 0.25 in the DMSO arm. Note that no relative AUC enrichment was observed between PDX models in the DMSO arm.

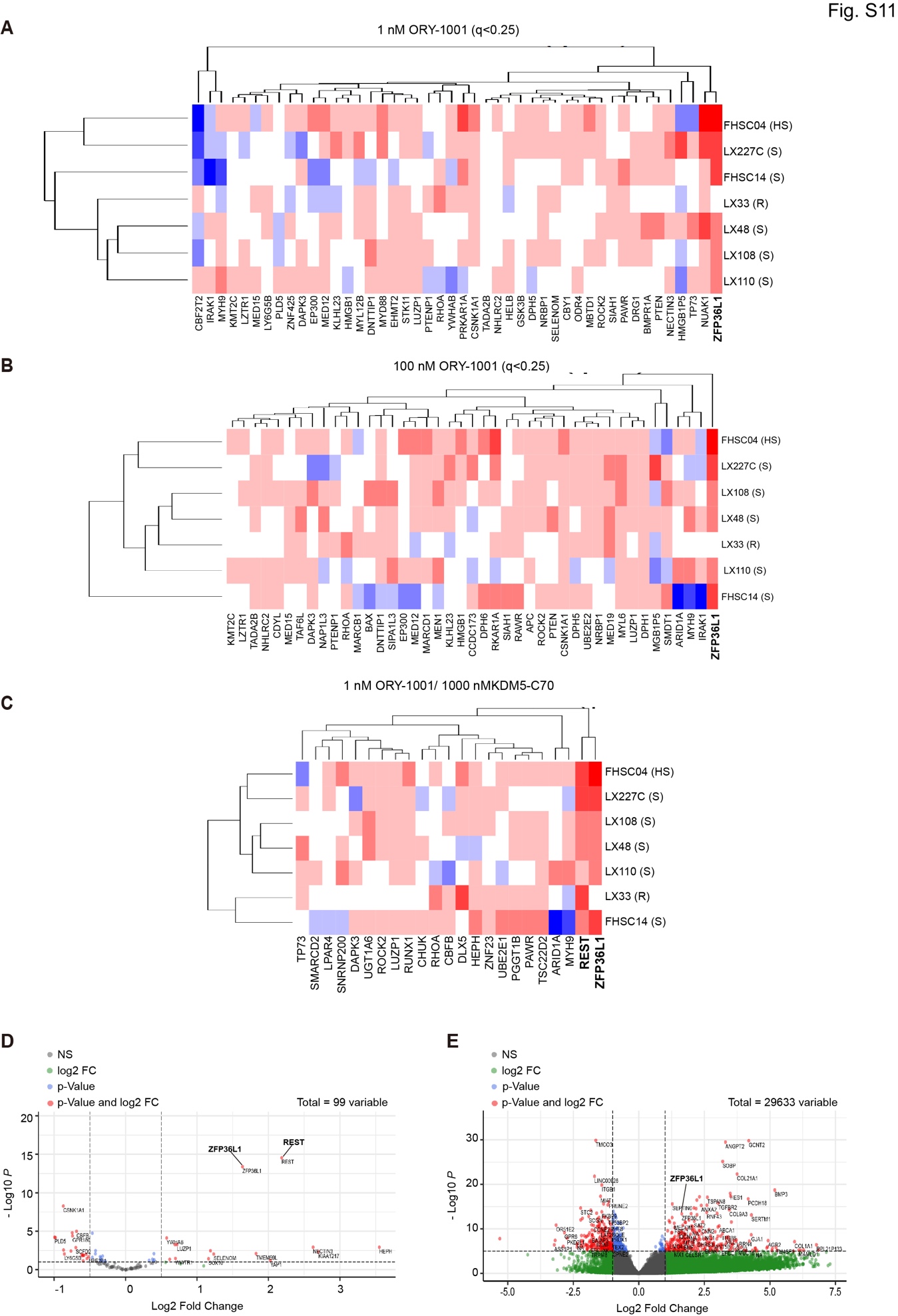

**Extended Data Fig. S11. Unsupervised Hierarchical Clustering of Hits from our CRISPR/Cas9 ORY-1001/KDM5-C70 Resistance Screen after ORY-1001 Treatment.**

(**A-C**) Unsupervised hierarchical clustering of hits from our ORY-1001 1 nM (**A**), ORY-1001 100 nM (**B**), and ORY-1001 (1 nM) + KDM5-C70 (1000 nM) (**C**) CRISPR/Cas9 resistance screen (see Figs. 1C-E) of RNA-seq data from SCLC PDX models treated ex-vivo with ORY-1001 relative to DMSO. Red denotes genes with high expression, and blue denotes genes with low expression. (**D and E**) Volcano plot of RNA-sequencing (RNA-seq) data from the highly sensitive SCLC PDX model FHSC04 treated with ORY-1001 or vehicle *in vivo* showing the log2 fold change and -Log10 p-value of gene expression of hits from our ORY-1001 (100 nM) + KDM5-C70 (1000 nM) CRISPR/Cas9 ORY-1001 resistance screen with q-values less than 0.25 (**D**) or all mRNAs (**E**). Note that similar to the *ex vivo* results observed in Fig. 6B and S11A-C, ZFP36L1 and REST are most statistically significantly induced *in vivo*.
